# Splice donor site sgRNAs enhance CRISPR/Cas9-mediated knockout efficiency

**DOI:** 10.1101/532820

**Authors:** Ignacio García-Tuñón, Verónica Alonso-Pérez, Elena Vuelta, Sandra Pérez-Ramos, María Herrero, Lucía Méndez, Jesús María Hernández-Sánchez, Marta Martín-Izquierdo, Raquel Saldaña, Julián Sevilla, Fermín Sánchez-Guijo, Jesús María Hernández-Rivas, Manuel Sánchez-Martín

## Abstract

CRISPR/Cas9 enables the generation of knockout cell lines and null zygotes by inducing site-specific double-stranded breaks. In most cases the DSB is repaired by non-homologous end joining, resulting in small nucleotide insertions or deletions that can be used to construct knockout alleles. However, these mutations do not produce the desired null result in all cases, but instead generate a similar, functionally active protein. This effect could limit the therapeutic efficiency of gene therapy strategies based on abrogating oncogene expression, and therefore needs to be considered carefully. If there is an acceptable degree of efficiency of CRISPR/Cas9 delivery to cells, the key step for success lies in the effectiveness of a specific sgRNA at knocking out the oncogene, when only one sgRNA can be used. This study shows that the null effect could be increased with an sgRNA targeting the splice donor site (SDS) of the chosen exon. Following this strategy, the generation of null alleles would be facilitated in two independent ways: the probability of producing a frameshift mutation and the probability of interrupting the canonical mechanism of pre-mRNA splicing. In these contexts, we propose to improve the loss-of-function yield driving the CRISPR system at the SDS of critical exons.

## INTRODUCTION

With the recent diversification of genome editing tools, including those involving clustered, regularly interspaced short palindromic repeats and their nuclease-associated protein Cas9 (CRISPR/Cas9), the landscape of suppression techniques has dramatically changed. Although CRISPR/Cas9 is similar in action and efficacy to protein-based targeted nucleases, such as zinc finger nucleases (ZFNs) and transcription activator-like effector nucleases (TALENs)(1), the ease with which these reagents can be designed and tested through the construction of single-guide RNAs (sgRNAs) has made gene editing available to a wider variety of users and for a broader range of applications. Unlike ribozymes, antisense oligodeoxynucleotides (AS-ODNs) and short interfering RNAs (siRNAs), CRISPR/Cas9 works at the DNA level, where it has the advantage of providing permanent and full gene knockout, while other methods only silence genes transiently(2)^,^(3). CRISPR/Cas9 cuts DNA in a sequence-specific manner with the possibility of interrupting coding sequences, thereby making it possible to turn off cancer drivers in a way that was not previously feasible in humans(4, 5). This notable application of permanent gene disruption is based on the cellular mechanisms involved in double-stranded break (DSB) repair. Following the creation of a DSB within the coding sequence of a gene, mechanisms of DNA repair can induce insertions and deletions (indels), resulting in frameshift or nonsense mutations(6). Its great efficiency at inducing DSB has led to CRISPR/Cas9 technology gaining a reputation as the gold standard for creating null alleles *in vivo* and *in vitro*. These null alleles can arise from frameshift mutations that trigger premature stop codons and/or non-sense-mediated decay in the target gene, resulting in loss of function. Currently, CRISPR/Cas9 is extensively used to engineer gene knockouts in most biological systems, but due to the variable size of the NHEJ-induced indel, it is not always possible to generate a full KO in one step. When the delivery of Cas9 elements is effective, full KO generation requires off-frame mutations in both alleles, which is a matter of probability since the random nature of DNA repair gives rise to considerable heterogeneity within the cell. It entails dealing with a significant frequency of mutated cells in which the outcome of mutation could preserve the reading frame (i.e., +3 or −3 mutations)(7). A possible solution is to use two or more RNA guides to knock out the gene at several key sites in an attempt to guarantee the null result. However, a high proportion of off-targets would increase with each new sgRNA added. Conversely, more sgRNAs at the same time trigger more DSBs, which induces a stronger p53-mediated DNA damage response(8) and more complex rearrangements(9). Either way, these undesirable effects may be irrelevant in assays in which the knockout cell can be sequenced, selected and expanded, or the null allele of the animal model can be segregated. Nevertheless, there are other situations, either *in vivo* or *in vitro*, in which cell selection and clone expansion are not available, and achieving high levels of knockout or gene inactivation efficiency is crucial(10, 11). Thus, it is important to study the key exons carefully and, more importantly, the target areas inside them, before making a selection(12). Hematological cancer therapies based on specific oncogenic silencing within primitive pluripotent stem cells may be the best example of these situations. In this pathological cell context, the highly efficient interruption of the oncogenic open reading frame (ORF) might be an effective therapeutic option. It would even be more important for those tumors directed by a single oncogenic event, as is the case for several leukemias or sarcomas, which are directed by specific fusion oncoproteins(13, 14). A recent study of the BCR/ABL oncogene showed this gene fusion to be an ideal target for CRISPR/Cas9-mediated gene therapy. A CRISPR-Cas9 application truncated the specific BCR-ABL fusion (p210) abrogating its oncogenic potential, but to achieve *in vivo* effectiveness in a xenograft model, the authors had to select and expand the correctly edited cellular clone because some of the clones contained in-frame or non-synonymous mutations(5, 15). Therefore, in these situations, it is essential to have not only highly efficient Cas9-sgRNA cell delivery, but also a high capacity for generating null mutations. This is especially critical for cancer oncogene suppression therapies based on disrupting driver oncogenes. If the efficiency of CRISPR/Cas9 reagent delivery to the cancer cell is acceptable, the key step to success lies in the effectiveness with which a specific sgRNA can knock out the oncogene. In this way, for most knockout studies in which the edited cells or mice can be selected, the sgRNA targets different positions within the chosen exon, avoiding exon boundaries. In most of these cases, the sgRNA design follows only off-target criteria, but for cases in which cellular selection is not an option and only one sgRNA can be used, the null effect could be strengthened with an sgRNA that targets splice site consensus sequences. Following this strategy, the generation of null alleles would be enhanced in two ways: by increasing the probabilities of producing a frameshift mutation and by breaking the canonical mechanism of pre-mRNA splicing. In this sense, it has long been known that mutations in splice-site consensus sequences can affect pre-mRNA splicing patterns and can lead to the generation of null or deficient alleles(16). In fact, pioneering genetic studies indicated that many of the thalassemia mutations in the β-globin gene affect splice sites and give rise to aberrant splicing patterns(17, 18). Recent studies have demonstrated that a splicing mutation in the STAR gene is a loss-of-function mutation that produces an aberrant protein(19). Nonsense-mediated mRNA decay (NMD), a conserved biological mechanism that degrades transcripts containing premature translation termination codons, could help secure the null effect when a DSB is induced at splice sites. In addition to transcripts derived from nonsense alleles, the substrates of the NMD pathway include pre-mRNAs that enter the cytoplasm with their introns intact(20). Several mutations of splice donor sites that cause loss of gene function have recently been identified. A novel mutation at a splice donor site that was predicted to lead to skipping of exon 10 of the PLA2G6 gene was found in a homozygous state in infantile neuroaxonal dystrophy patients. This variant was correlated with very strong loss of function, providing further evidence of its pathogenicity(21). Mutations in the ectodysplasin A1 gene (EDA-A1) at the splice donor site have been described in patients with hypohydrotic ectodermal dysplasia. This novel functional skipping-splicing EDA mutation was the cause of the pathological phenotype(22). Studies in a family with premature ovarian failure identified a variant that alters a splice donor site. This variant resulted in a predicted loss of function of the MCM9 gene, which is involved in homologous recombination and repair of double-stranded DNA breaks(23).

Taking into account all these findings, we decided to explore the effectiveness of driving one single sgRNA targeting the splice-donor exon site (SDE-sgRNA) to increase the null allele yield. To compare the knockout efficiency of SDE-sgRNAs and sgRNAs targeting positions within the exon (IE-sgRNA) we induced DSB with both guides in critical exons in three genes (TYR, ATM and ABL), two systems (*in vivo* and *in vitro)*, and two species (human and mouse). Finally, we sequenced all mutant alleles generated and analyzed the consequences *in silico* and *in vivo*.

## MATERIAL & METHODS

### Cell lines and culture conditions

Baf/3 is a murine interleukin 3-dependent murine pro-B cell(24). Baf/3was maintained in Dulbecco’s Modified Eagle’s Medium (DMEM) (Life Technologies) supplemented with 10% fetal bovine serum (FBS) and 1% of penicillin/streptomycin (Life Technologies) and 10% of WEHI-3-conditioned medium, as a source of IL-3.

The human CML-derived cell lines K562 were purchased from Deutsche Sammlung von Mikroorganismen and Zellkulturen (DMSZ). K562 cells were cultured in RPMI 1640 medium (Life Technologies) supplemented with 10% FBS, and 1% penicillin/streptomycin (Life Technologies). All cell lines were incubated at 37ºC in a 5% CO_2_ atmosphere. The presence of mycoplasma was tested frequently in all cell lines with a MycoAlert kit (Lonza), using only mycoplasma-free cells in all the experiments carried out.

### CRISPR/Cas9 system design and sgRNA cloning

pX458 (Addgene plasmid # 48138)(25), which contains the coding sequence of Cas9 nuclease and GFP, and a cloning site for sgRNA sequence, was digested with BpiI (NEB). To clone the sgRNAs into the pX458 vector, two complementary oligos were designed for each sgRNA that included two 4-bp overhang sequences (Table 9). The sgRNA sequences were designed with the web tool of the Spanish National Biotechnology Centre (CNB)-CSIC (http://bioinfogp.cnb.csic.es/tools/breakingcas/).

**Table 9.**
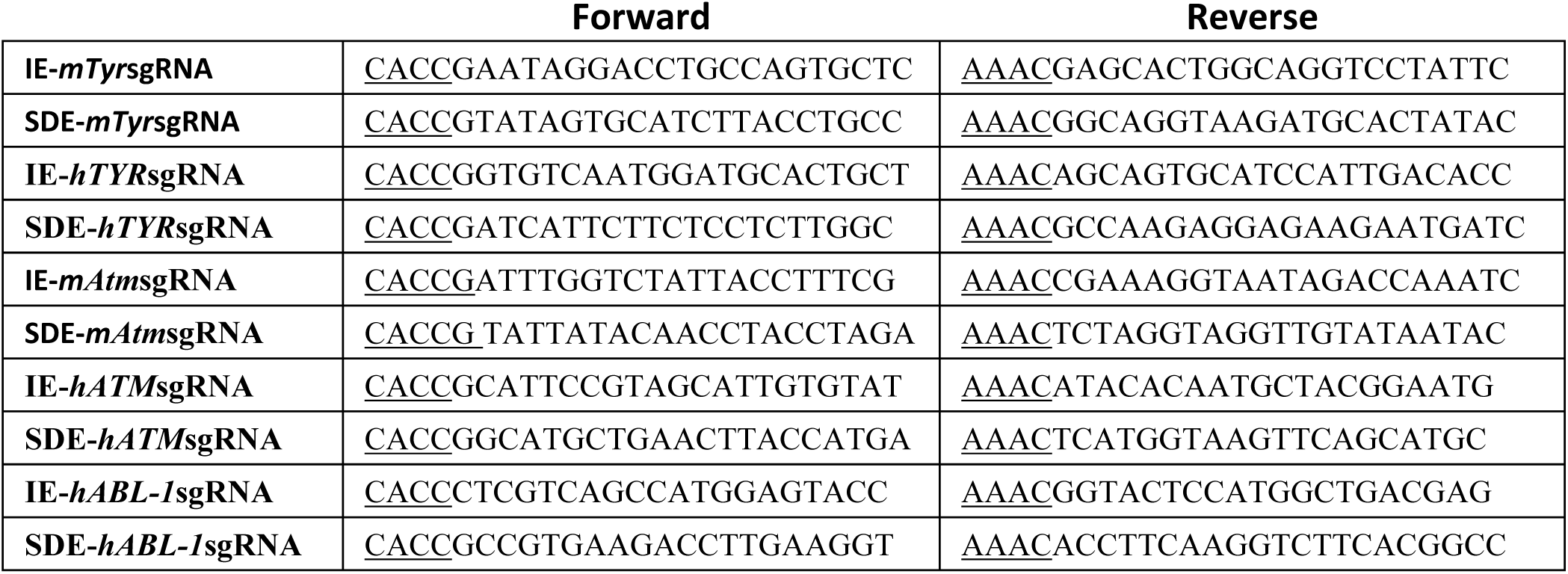
Oligos designed for each sgRNA.

Two sgRNAs were designed for the *Tyr* mouse locus. One of them, IE-*mTyr*sgRNA, targets the exonic sequence in *Tyr* exon1, and the other, SDE-*mTyr*sgRNA, targets the exon1-intron1-2 junction (Figure 1A). Two sgRNAs were designed to target homologous sequences in the human *TYR* locus: IE-*hTYR*sgRNA and SDE-*hTYR*sgRNA (Figure 3A).

**Figure 1.**
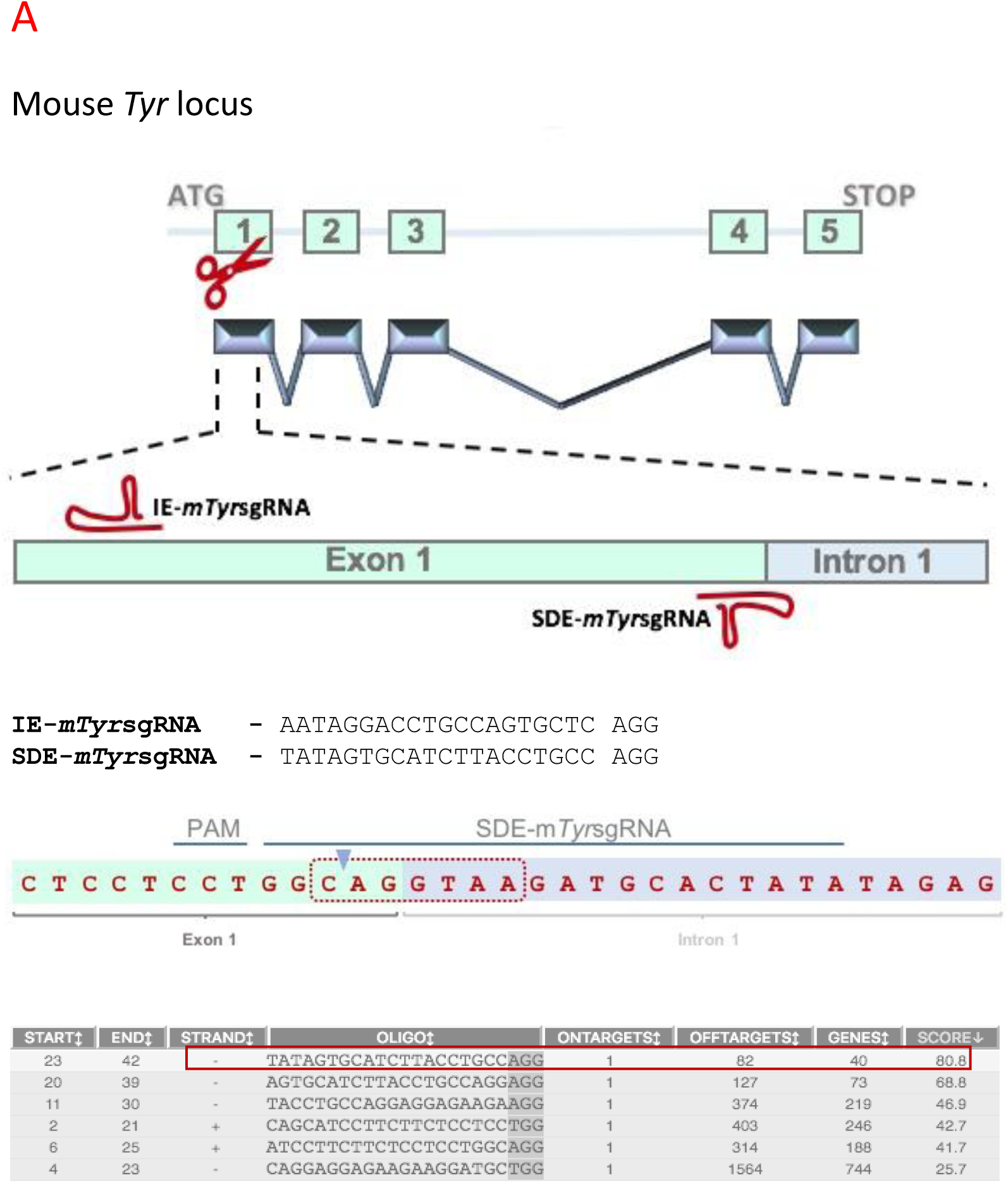
CRISPR/Cas9 edition of mouse *Tyr* locus *in vitro*. (A) **Schematic representation of the mouse *Tyr* locus and the CRISPR/Cas9 experimental design the two RNA guides are represented in the exon 1 sequence.** SDE-*mTyr*sgRNA is complementary to the splice site between exon 1 and intron 1-2. IE-*mTyr*sgRNA targeted a central position at the coding sequence of exon1. SgRNA sequences and their respective PAM sequences are also described. Exon-intron junction and nucleotides implicated in intron processing (red dot line). SDE-sgRNA (red box) was chosen among several candidates based on its score.

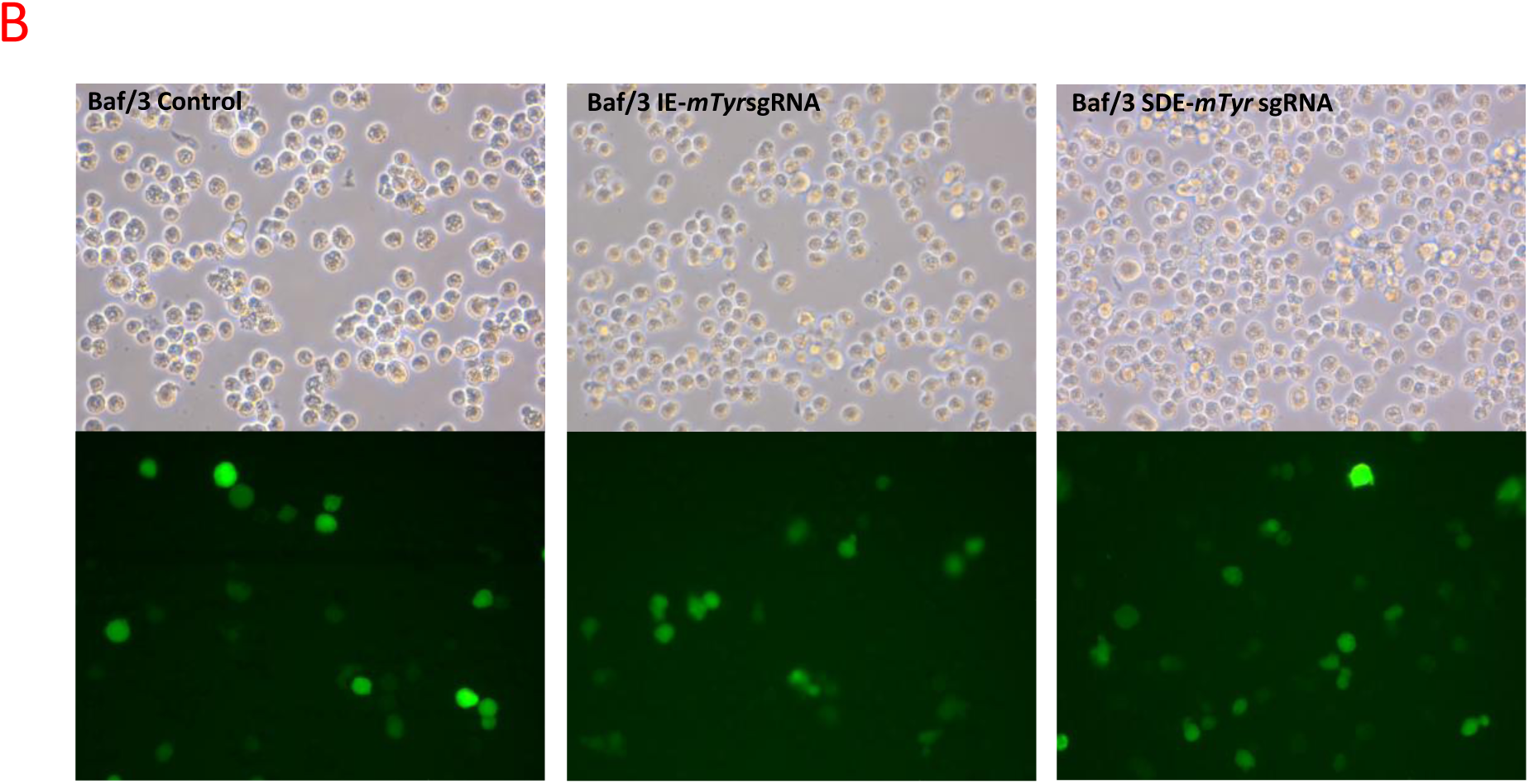
*(B) Tyr* locus CRISPR/Cas9-mediated edition in the BOFF p210 mouse cell line. Fluorescent microscopy of Boff cells electroporated with empty px480 vector and carrying each RNA guides. (C) CRISPR/Cas9 edited *Tyr* sequences of Boff cells through IE-*mTyr*sgRNA (red box) and SDE-*mTyr*sgRNA (blue box). Control cells showed a wt sequence of *Tyr* gene, while Boff edited cells showed a mixture of sequences around the expected cleavage point for each sgRNA. TIDE analysis of sequences predicted the overall edition efficacy and most common allele variations generated.

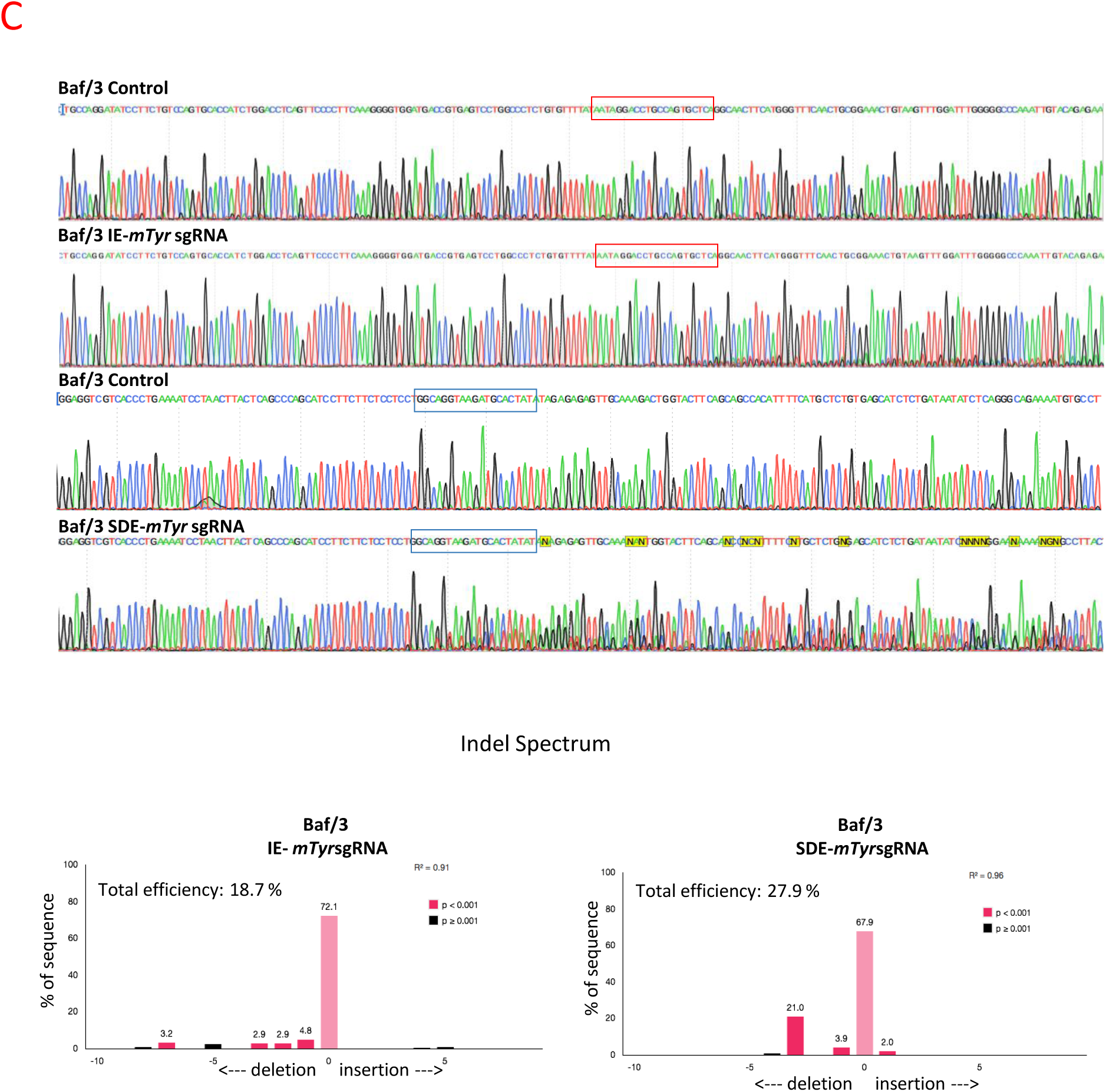
(C) CRISPR/Cas9 edited *Tyr* sequences of Boff cells through IE-*mTyr*sgRNA (red box) and SDE-*mTyr*sgRNA (blue box). Control cells showed a wt sequence of *Tyr* gene, while Boff edited cells showed a mixture of sequences around the expected cleavage point for each sgRNA. TIDE analysis of sequences predicted the overall edition efficacy and most common allele variations generated.

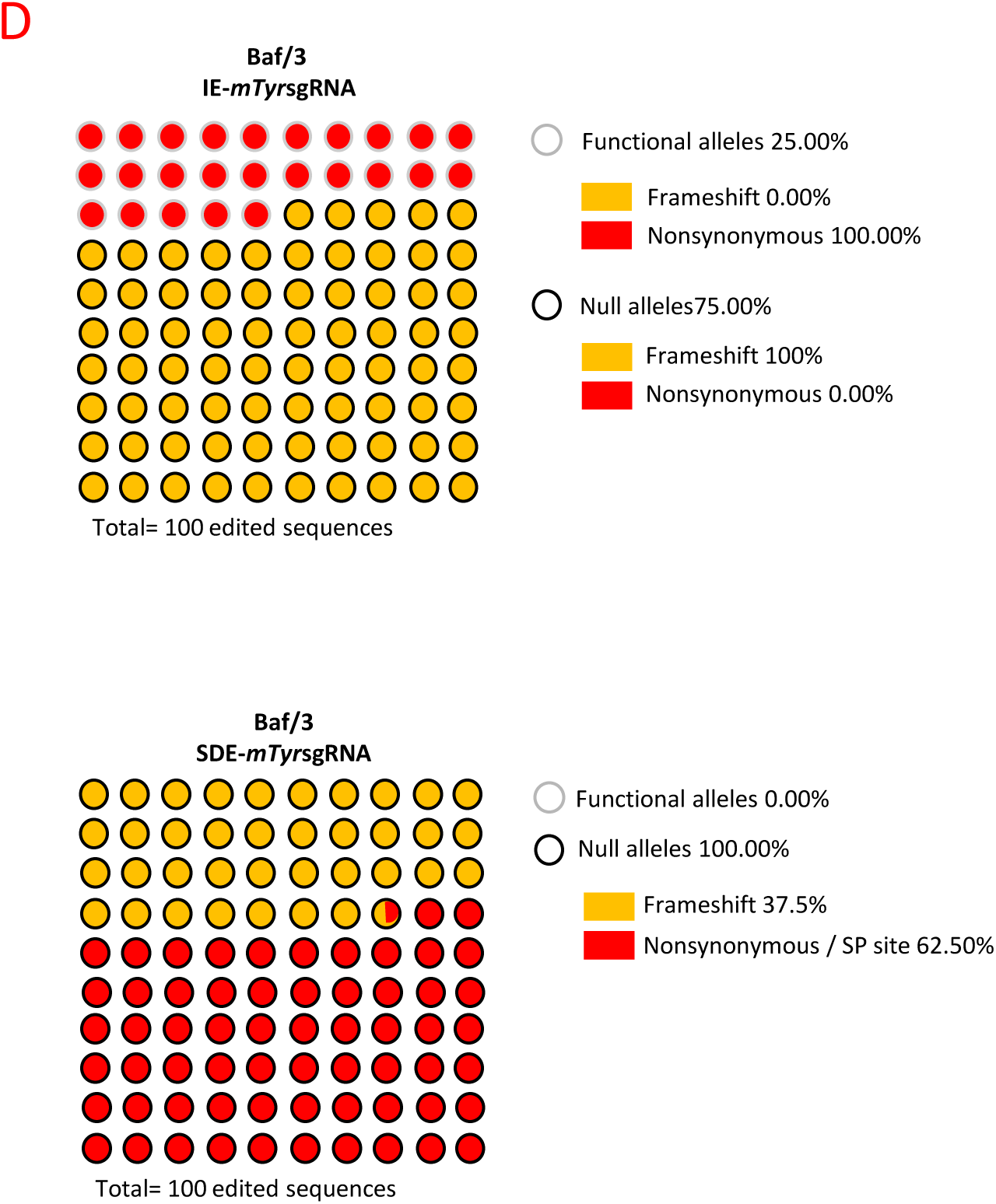
(D) Next Generation Sequencing (NGS) of *Tyr* gene-edited Boff cells. Graphic representation of the mutations found, and their predicted effect in cells edited by IE-*mTyr*sgRNA and SDE-*mTyr*sgRNA. Black and gray circles represent null alleles and functional alleles respectively, while the background indicates the type of mutation (red: nonsynonymous/sp site; yellow: frameshift). Only 75% of IE-*mTyr*sgRNA-edited sequences resulted in a null allele, while 100% of SDE-*mTyr*sgRNA-edited sequences gave rise to a loss of gene function.

In the same way, two sgRNAs against the mouse *Atm* locus (IE-*mAtm*sgRNA and SDE-*mAtm*sgRNA; Figure 5A) and two sgRNAs against the human *ATM* locus (IE-*hATM*sgRNA and SDE-*hATM*sgRNA; Figure 4A) were designed, one of each pair in the coding sequence of exon 10 (IE) and the other against the *ATM* exon10-intron10-11 splice donor exon (SDE).

Finally, two sgRNA against human *ABL-1* locus were designed: IE-*hABL-1*sgRNA, which targets the exon 6 coding sequence, and SDE-*hABL-1*sgRNA, which targets the exon 4 splice donor sequence.

The two complementary oligos used to conform each sgRNA (Table 9) were denatured at 95ºC for 5 min, ramp-cooled to 25ºC over 45 min to allow annealing, and finally ligated with the linearized px458. 2 μl of the ligation reaction were used to transform competent cells, and single colonies were expanded using a QIAprep spin Maxiprep Kit (Qiagen) before plasmid extraction. The correct insertion of the sgRNA sequences was confirmed by Sanger sequencing.

### *In vitro* cell electroporation

Mouse Baf/3Baf/3and human K562 cells were electroporated with px458 containing sgRNAs against the *Tyr* and *ATM* loci, respectively, using Amaxa Nucleofector II (Lonza). 2 x 10^6^Boff p210 mouse cells were electroporated with 15 μg of plasmid in 100 μl of electroporation buffer (5 mM KCl; 15 mM MgCl_2_; 120 mM Na_2_HPO_4_/NaH_2_PO_4_pH7.2; 25 mM sodium succinate; 25 mM manitol)(26) using program X001, while 1 x 10^6^k562 cells were electroporated with 10 μg of plasmid using program T016. 24 hours after electroporation, GFP-positive cells were sorted by fluorescence-activated cell sorting (FACS) using FACS-Aria (BD Bioscience). 72 hours post-electroporation, the genome editing of the cells was analyzed.

### Sequencing of sgRNA targets sites

Genomic DNA was extracted using the QIAamp DNA Micro Kit (Qiagen) following the manufacturer’s protocol. To amplify the different target regions of human and mouse *TYR* and *ATM* genes, and human *ABL-1*, PCR was performed with the oligos described in Table 3. Genomic DNA from single blastocyst-staged embryo was extracted in 10 μl of lysis buffer (50 mM KCL, 10 mM Tris-HCL pH 8.5, 0.1% Triton x-100, and 4 mg/ml of proteinase K) at 55ºC overnight, then heated at 95ºC for 10 min. 2 μl of this DNA solution was used as a template for two rounds of PCR (30 cycles + 20 cycles) to amplify the target sequences using a specific primer for each region (Table 11).

**Table 3.**
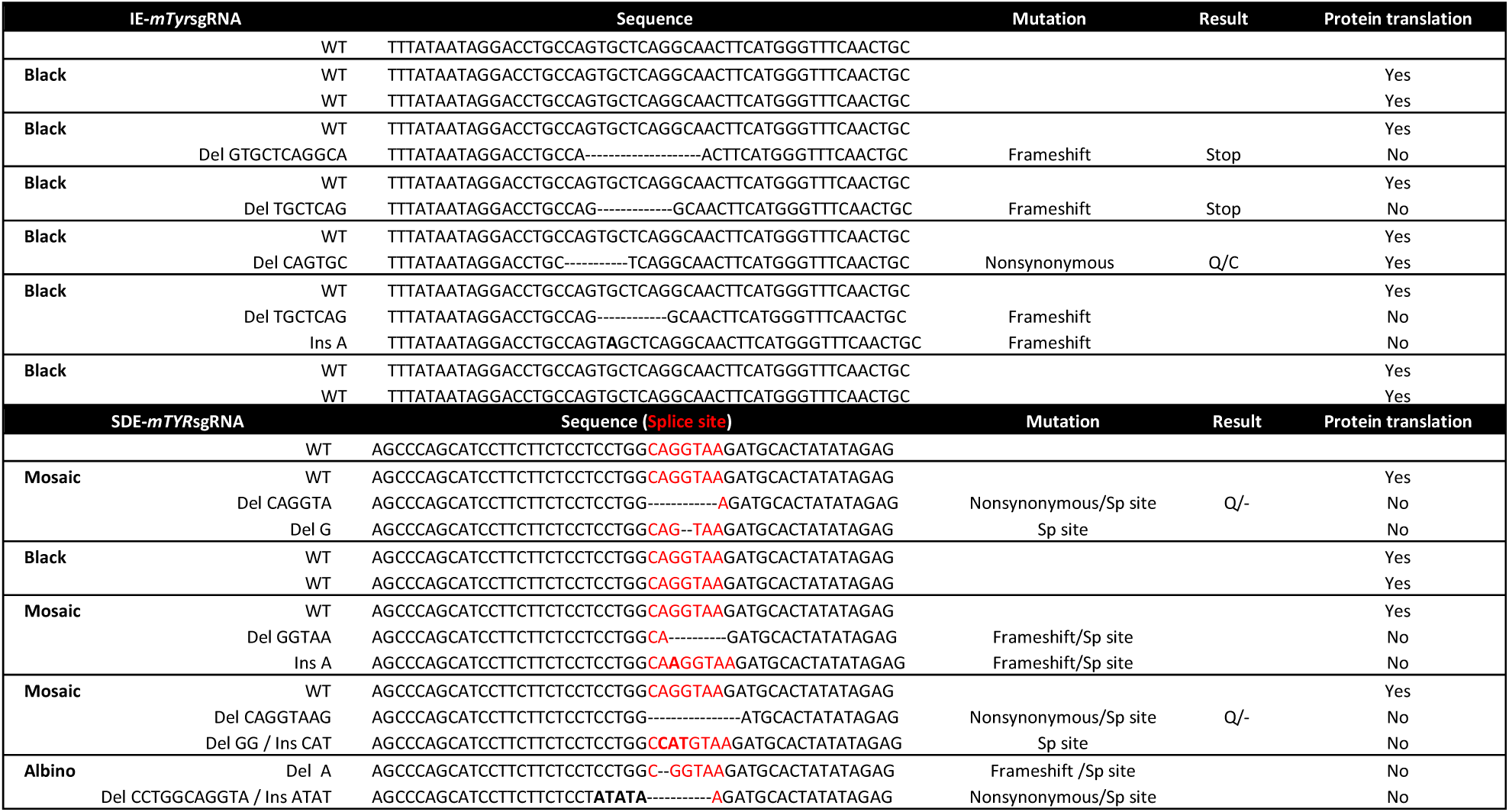
*In vivo* genome editing of *Tyr* locus in mice using sgRNA against the coding sequence (IE) and the coding SDE sequence. Observed phenotype and Sanger analysis of allelic variants induced in mice born after CRISPR/Cas9 system microinjection.

**Table 11.**
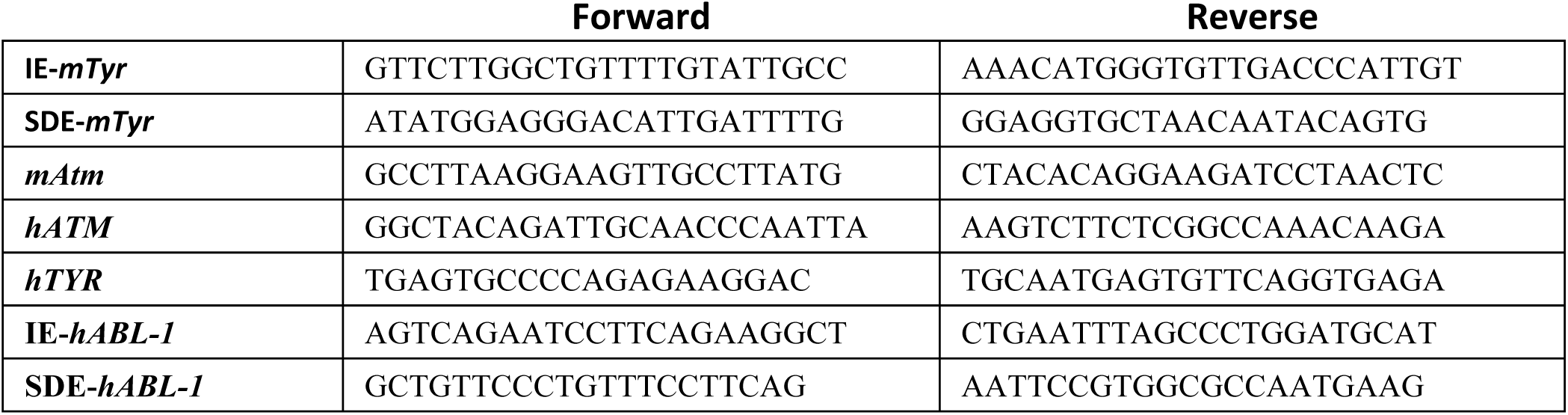
Oligos used for target genome sequence amplification.

PCR products were purified using a High Pure PCR Product Purification Kit (Roche) and sequenced by the Sanger method using forward and reverse PCR primers.

The editing efficiency of the sgRNAs and the mutations potentially induced were assessed using Tracking of Indels by Decomposition (TIDE) software (https://tide-calculator.nki.nl; Netherlands Cancer Institute), which only required two Sanger sequencing runs from wild-type cells and mutated cells.

To specifically identify the different generated mutations, Next Generation Sequencing (NGS) technology was employed with the same Sanger primers with the corresponding adapters added, to read each edited sequence individually.

The purified amplicons were mixed in equimolar ratios according to the number of molecules and diluted to a final concentration of 0.2 ng/ul. The indexed paired-end library was prepared with a Nextera XT DNA Sample Preparation Kit (Illumina) and sequenced using an Illumina platform (NextSeq or MiSeq, 300 cycles). A median per base coverage of 27,538 reads (range 2096–88,976) was achieved. To call the sequence variants, an in-house bioinformatics pipeline was established. Sequencing reads were aligned to the mouse reference sequence genome (mm9) using bwa-0.7.12 software, and variant calling was performed with VarScan.v2.4. To visualize read alignment and confirm the variant calls, Integrative Genomics Viewer version 2.3.26 (IGV, Broad Institute, MA) was used.

### Flow cytometry analysis and cell sorting of single-edited cell-derived clone

72 hours after sgRNA electroporation of K562 and Boffp210 cells, GFP-positive cells were selected by fluorescence-activated cell sorting (FACS) using FACS Aria (BD Biosciences), establishing the edited K562 and Boff cell pool lines. For K562, single cells were seeded in 96-well plates by FACS, establishing six random single-cell-derived clones for both *ATM* sgRNAs, and used to analyze ATM protein expression. Six clones derived from cells electroporated with empty vector were used as controls.

### Western blotting

ATM protein expression was assessed by SDS-PAGE and western blot using a rabbit anti-ATM antibody (1:1000; 2873S; Cell Signaling). Horseradish peroxidase-conjugated α-rabbit antibody (1:5000; 7074S; Cell Signaling) was used as a secondary antibody. Antibodies were detected using ECL^™^Western Blotting Detection Reagents (RPN2209, GE Healthcare). The expression of vinculin (rabbit anti-vinculin; 1:1000; 4650S; Cell Signaling) was used as a loading control.

### *In vitro* transcription of CRISPR/Cas9 system components, animals and embryo microinjection

Both *Tyr* sgRNA sequences were PCR-amplified from px458-based vector with primers carrying the T7 RNA polymerase promoter at the 5′ ends (Table 10), and after column purification (Roche) the resulting PCR was used as a template for T7 RNA polymerase transcription *in vitro* (MEGAshortscript™ T7 Transcription Kit, Thermo Fisher).

**Table 10.**
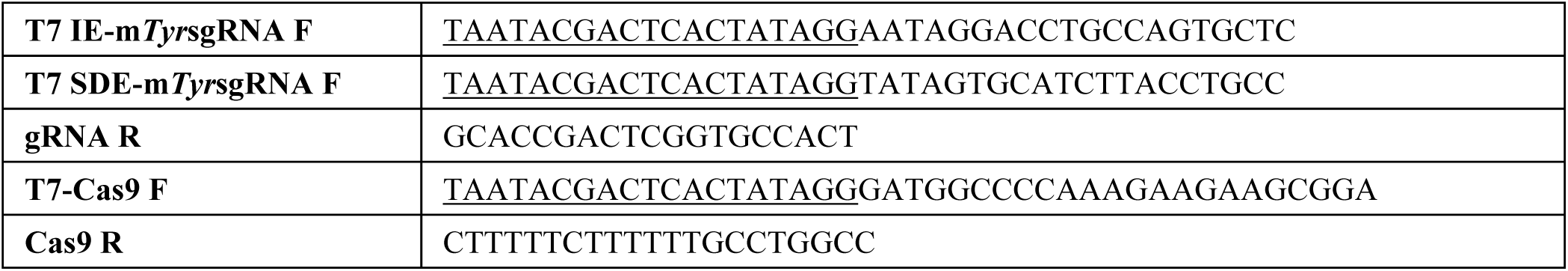
Oligos used for *in vitro* transcription of sgRNA and Cas9 mRNA.

The Cas9 nuclease ORF, including NLS, was also PCR-amplified with primers carrying the T7 RNA polymerase promoter at the 5′ ends (Table 10). The PCR product was purified and used as a template for *in vitro* transcription, 5′ capping (mMESSAGE mMACHINE™ T7 Transcription Kit, Thermo Fisher), and 3′ poly(A) tailing (Poly(A) Tailing Kit, Thermo Fisher). Transcription products were purified with RNeasy Mini Kit (Qiagen) and eluted in nuclease-free EmbryoMax microinjection buffer (Millipore).

One-cell-staged embryos from superovulated C57BL/6J or B6/CBA hybrid females were harvested and microinjected with 20 ng/μl of sgRNA and 20 ng/μl of Cas9 mRNA into the cytoplasm and pronucleus. This study followed Spanish and European Union guidelines for animal experimentation.

### Statistical analysis

Statistical analysis of annexin V expression was performed using GraphPad Prism version 6.00 for Mac OS X, (GraphPad Software, La Jolla California USA, www.graphpad.com). Experimental results were expressed as median ± standard error (SEM). Nonparametric variables were analyzed using Kruskal-Wallis followed by Dunn′s multiple comparisons test. Values with p<0.001 (indicated by three asterisks) were considered to be statistically significant.

## RESULTS

### 1. The sgRNA guide targeting the splice-donor site of mouse Tyr exon 1 increases the generation of null alleles in *in vitro* and *in vivo* systems

Two sgRNAs were created to study the efficiency of SDE-sgRNA and IE-sgRNA at generating null alleles in mouse cells (Figure 1A). Both guides were designed to target the tyrosinase mouse gene at exon 1 in a pro-B mouse cell line (Baf/3) and C57Bl/6J mouse zygotes. Previously, we sequenced Tyr exon1 and no nucleotide polymorphisms at the sgRNA target sites were found in Baf/3 cells or C57Bl/6J zygotes.

#### 1.1. *In vitro* assays in the Baf/3 mouse cell line

Three individual electroporation assays of Baf/3 cells were performed, with each sgRNA directed to Tyr exon 1 (SDE-sgRNA and IE-sgRNA) cloned in a CRISPR-Cas9-GFP mammalian expression vector. An empty CRISPR-Cas9-GFP vector was used as a control. GFP expression was detectable 24 hours post-electroporation in all cases, indicating the effective delivery of the CRISPR/Cas9 system and its expression in Baf/3 cells (Figure 1B).

Sanger sequencing identified indel mutations at the predicted cleavage point in CRISPR/Cas9 assays, while no sequence variations were observed in control cells (Figure 1C). Tracking of indels by decomposition (TIDE) analysis showed similar overall DSB-induced efficiency for SDE-sgRNA and IE-sgRNA in the Baf/3 cell line. To eliminate interference in Cas9 delivery efficiency among assays, we decided to analyze only the mutant alleles generated by both guides and their consequences for the obviation of wildtype or well-repaired alleles.

In knockout assays with both sgRNAs, the TIDE algorithm of Baf/3 mutant cells predicted small deletions (1-4 bp) in most cases: 76.2% with IE-*mTyr*sgRNA (R^2^= 0.91, p < 0.001) and 85% with SDE-*mTyr*sgRNA (R^2^= 0.96, p < 0.001) (Figure 1C).

To gain detailed information about all mutant alleles for each sgRNA we analyzed the genome of properly electroporated Baf/3 cells by next-generation sequencing (NGS) (Table 1). Unlike with the Sanger analysis, NGS revealed a high number of mutated alleles: 17 with IE-*mTyr*sgRNA and nine with SDE-*mTyr*sgRNA. NGS revealed mutated alleles in the IE-*mTyr*sgRNA and SDE-*mTyr*sgRNA groups with nonsynonymous indels that deleted two or six amino acids, thereby preserving the reading frame of the protein (Figure 1D, Table 1). However, *in silico* analysis of the allelic modifications generated by SDE-*mTyr*sgRNA predict the generation of a null allele in all cases, by frameshift mutations or by loss of canonical splicing sequences, or both simultaneously (Figure 1D, Table 1).

**Table 1.**
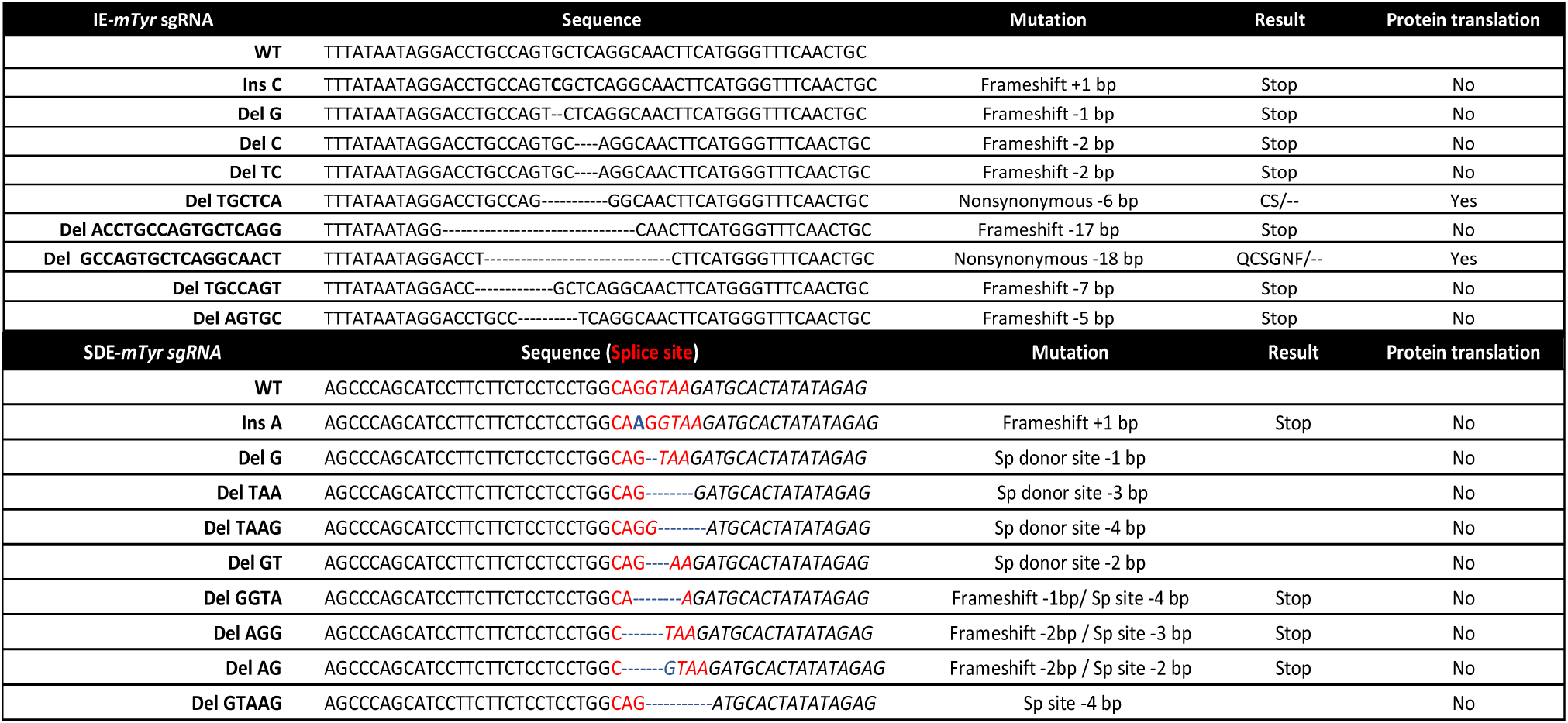
*In vitro* genome editing of the mouse *Tyr* locus using sgRNA against exon coding sequence (IE) and the coding splice-donor exon (SDE) sequence. NGS analysis of allelic variants induced in Baf/3 mouse cells.

#### 1.2. *In vivo* assays in mouse zygotes

One-cell stage embryos from two strains of mice, inbred C57Bl6/J and F2 hybrids of B6/CBA, were microinjected with Cas9 mRNA and Tyr sgRNAs. No nucleotide polymorphisms between C57Bl6/J and CBA strains at Tyr exon1/intron1 were found. The microinjected embryos were divided in two groups, one of which was grown to blast stage and harvested to obtain the genomic DNA, which was analyzed to detect indels at the sgRNA cut-sites. Embryos of the other group were grown to the two-cell stage and implanted in pseudo-pregnant females to visualize the *in vivo* CRISPR effect on mouse coat color.

##### 1.2.1. *In silico* analysis predicts a stronger knockout effect with sgRNA guides targeting the splice-donor site of mouse Tyr exon 1

The microinjected zygotes grown to blast stage were harvested to obtain their genomic DNA, which was then analyzed by NGS, revealing a greater abundance of null alleles in the SDE-*mTyr*sgRNA than in the IE-*mTyr*sgRNA embryo group (100% *vs.* 57.14%) (Table 2). Briefly, NGS detected seven mutated alleles at the expected cut-site of IE-*mTyr*sgRNA. *In silico* analysis identified three mutated alleles with nonsynonymous mutations that gave rise to a putative functional protein. NGS in the group of embryos microinjected with SDE-*mTyr*sgRNA identified eight mutated alleles, of which two were nonsynonymous mutations and six were null mutations. However, in this embryo group, all alleles (100%) detected were predicted to be null alleles given the splicing site mutations (Figure 2A, Table 2).

**Figure 2.**
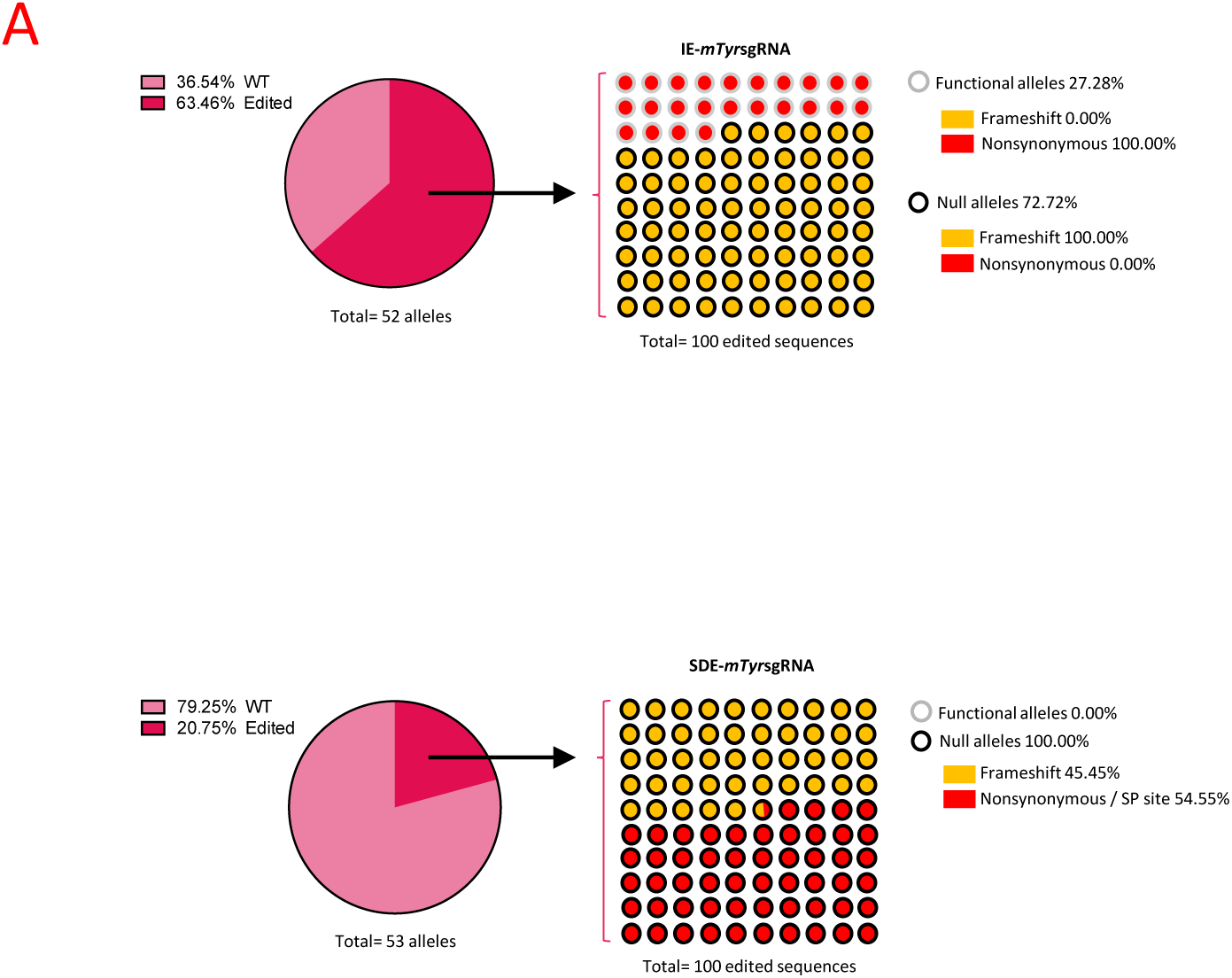
CRISPR/Cas9 editing of mouse *Tyr* locus *in vivo*. (A) NGS analysis of CRISPR/Cas9-mediated edition of *Tyr* locus in mouse blastocyst. Genotyping of embryos microinjected with sgRNAs against *Tyr* gene, by Next Generation Sequencing, revealed that only 27.28% of edited sequences from embryos microinjected with IE-m*Tyr*sgRNA correspond to functional alleles, while all SDE-*mTyr*sgRNA-modified alleles gave rise to loss-of-function alleles. Black and gray circles correspond to null and functional alleles, respectively, while the background indicates the type of mutation (red: nonsynonymous/sp site; yellow: frameshift).

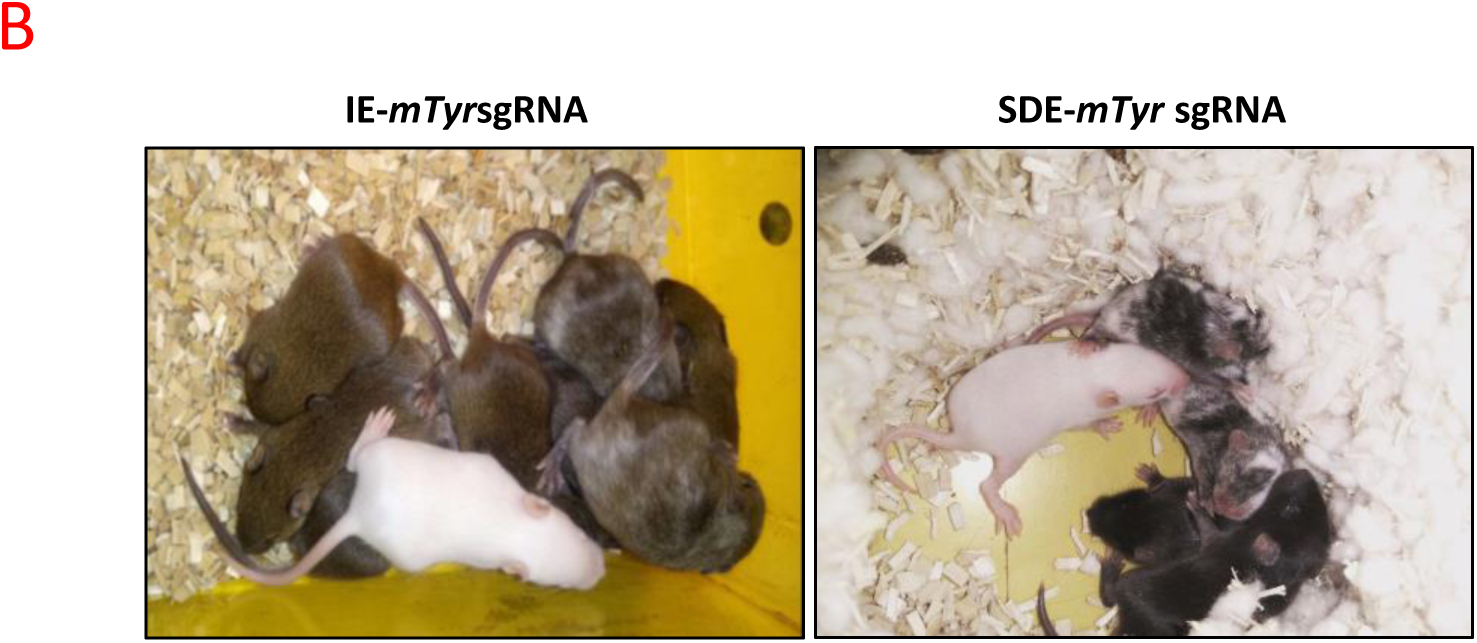
(B) Representative off-spring obtained from IE-*mTyr*sgRNA and SDE-m*Tyr*sgRNA. Most pups of IE-*mTyr*sgRNA-edited embryos (5 of 6) had a wt phenotype (black), while SDE-*mTyr*sgRNA gave rise to 1 wt, 1 albino and 3 mosaic pups.

**Table 2.**
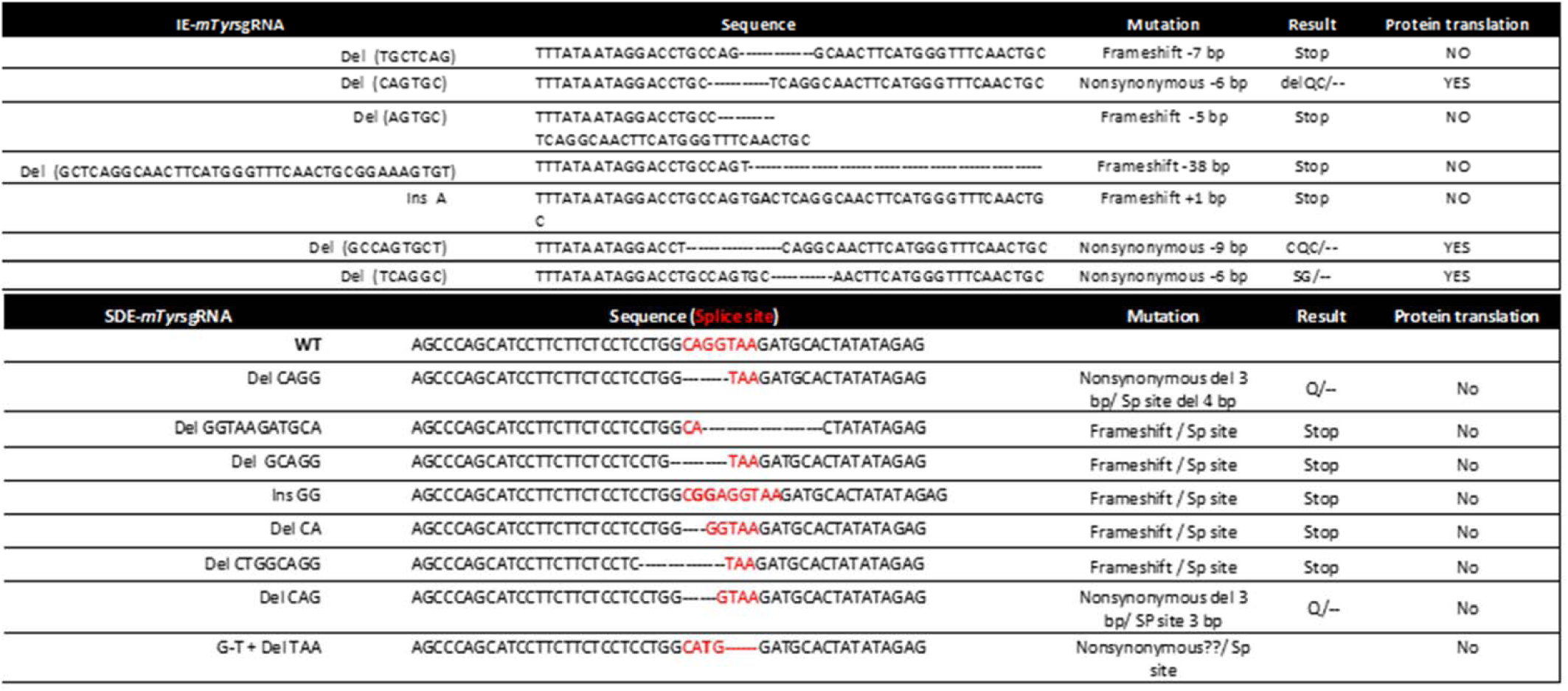
*In vivo* genome editing of *Tyr* locus in mouse embryos using sgRNA against the coding sequence (IE) and the SDE sequence. NGS analysis of allelic variants induced in microinjected mouse blastocysts.

##### 1.2.2. Functional analysis

To confirm the *in silico* predictions, one-cell stage embryos from two strains of mice were microinjected with Cas9 mRNA and both Tyr sgRNAs separately. Embryos microinjected with SDE-*mTyr*sgRNA or IE-*mTyr*sgRNA were implanted in two cell-stage in pseudopregnant females. Full albinos, mosaics, and colored-coat pups were detected in all offspring of each group of microinjected embryos in both strains (Figure 2B). 60 mice per group were analyzed by Sanger sequencing and a large number of mutant mice with one or two mutant alleles were detected. To address which sgRNA yielded a higher proportion of null alleles, we excluded all mice with unmuted alleles. All mice with at least one mutant allele (mosaic mice) were analyzed *in silico*. We detected a higher number of albino or mosaic mice in the SDE-*mTyr*sgRNA mouse group compared with the IE-*mTyr*sgRNA group (Table 3).

Sanger sequencing and TIDE analysis of the SDE-*mTyr*sgRNA mouse group with any grade of albinism identified at least two alleles with frameshift mutations and/or splice mutations. We detected full knockout cells in mosaic mice with three alleles: a wildtype allele, a frameshift null allele and a splicing-site-mutated allele arising from a point mutation (+1 bp insertion) at the intronic splice-site (Figure 2B and Table 3). We also detected coat-colored pups in IE-*mTyr*sgRNA targeted pups exclusively with two mutated alleles: a frameshift allele and a mutated allele arising from a nonsynonymous mutation (Table 3).

### 2. The sgRNA guides targeting the exon splice donor site enhances the knockout efficiency in human K562 cells

To ascertain whether the effects observed in mouse cells were specific to *Mus musculus*, two analog sgRNAs targeting human TYR exon 1 sequence were designed (IE-*hTYR*sgRNA and SDE-*hTYR*sgRNA) to target the leukemic BCR/ABL human K562 cell line (Figure 3A). Three individual electroporation assays in K562 cells were performed with each of the guides. GFP expression was detectable in all cases 24 hours post-electroporation, indicating an effective delivery and expression of CRISPR/Cas9 reagents in K562 cells (Figure 3B). GFP+ cells were sorted and subjected to Sanger sequencing, which revealed no variations in the target sequence of control cells (Figure 3B). As expected, we found indel mutations at the predicted cleavage point of IE-*hTYR*sgRNA and SDE-*hTYR*sgRNA. In the same way as in mouse BaF/3 cells, these alterations mainly corresponded to small indels (5-bp deletions and 1-bp insertions for IE-*hTYR*sgRNA, and 1-2-bp deletions and 1-bp insertions for SDE-*hTYR*sgRNA) (Figure 3C).

**Figure 3.**
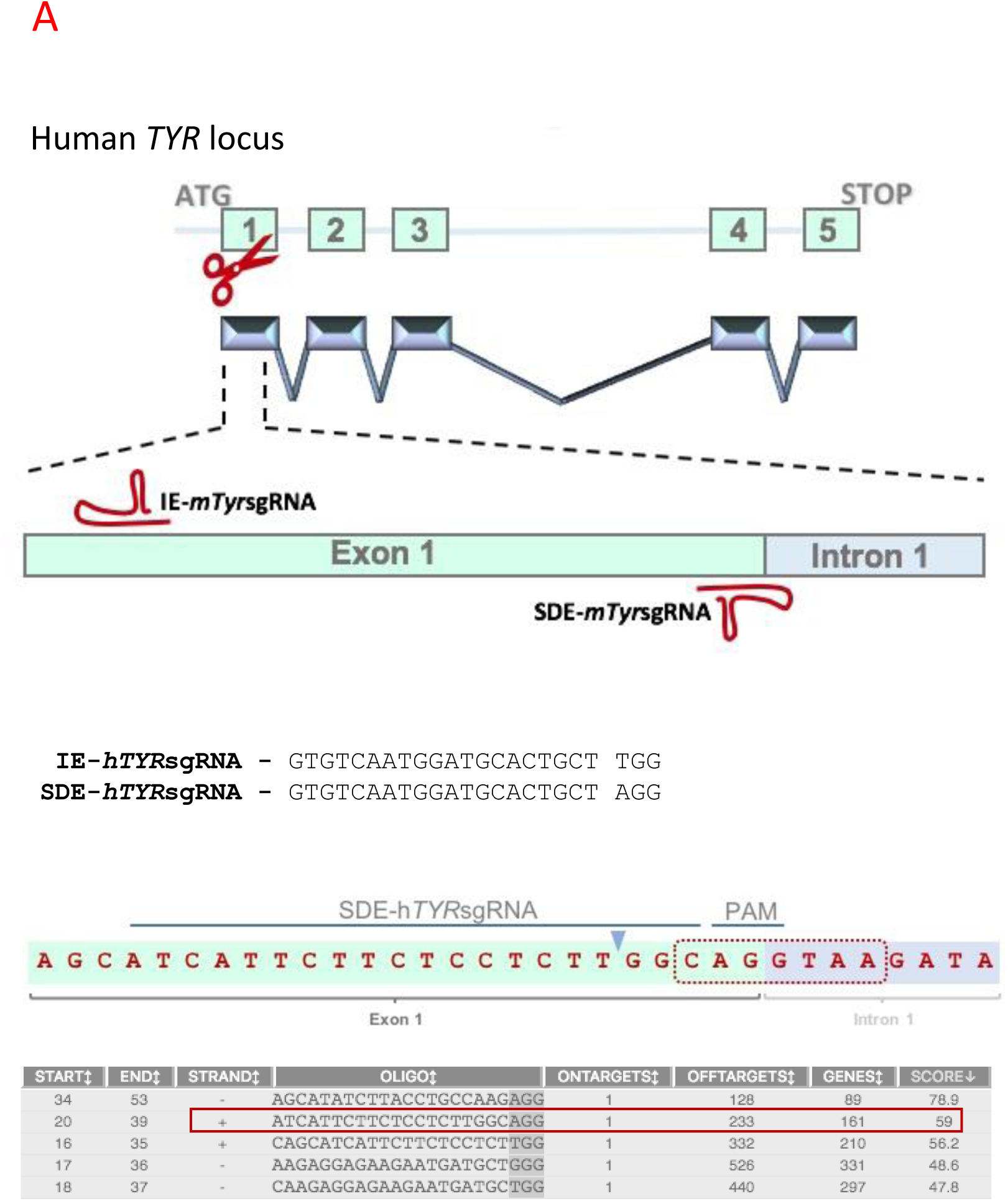
CRISPR/Cas9 editing of human *TYR* locus. (A) **Schematic representation of the human *TYR* locus and the CRISPR/Cas9 experimental design in which the two RNA guides are represented in the exon 1 sequence (red).** SDE-*hTYR*sgRNA is complementary to the coding sequence of the splice site between exon 1 and intron 1-2. IE-*hTYR*sgRNA targeted a central position at the coding sequence of exon 1. sgRNA sequences and their respective PAM sequences are also described. Exon-intron junction and nucleotides implicated in intron processing (red dot line). SDE-sgRNA (red box) was chosen among several candidates based on its score.

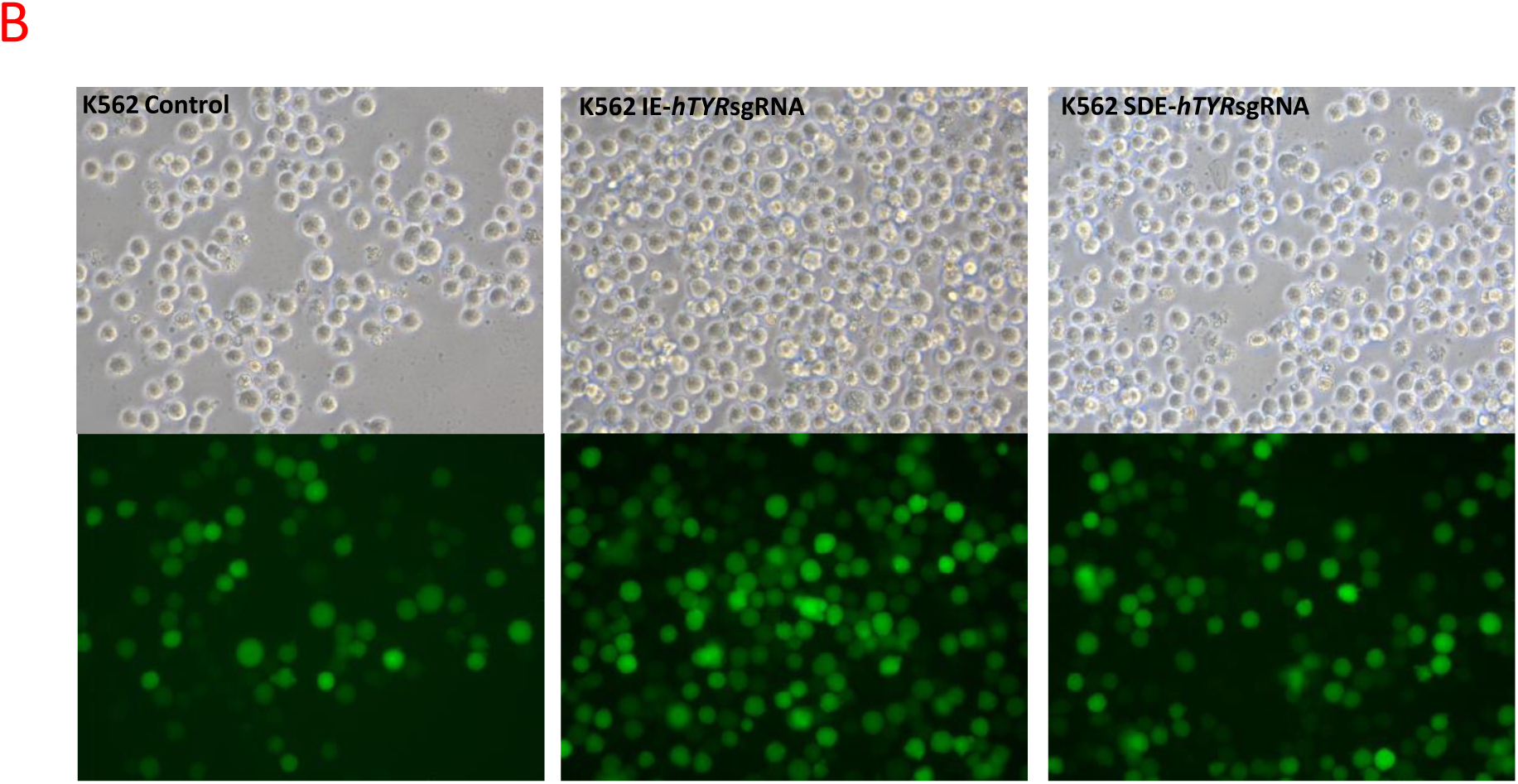
*(B) TYR* locus CRISPR/Cas9-mediated editing in the K562 human cell line. Fluorescent microscopy of K562 cells electroporated with empty px480 vector and carrying each RNA guide.

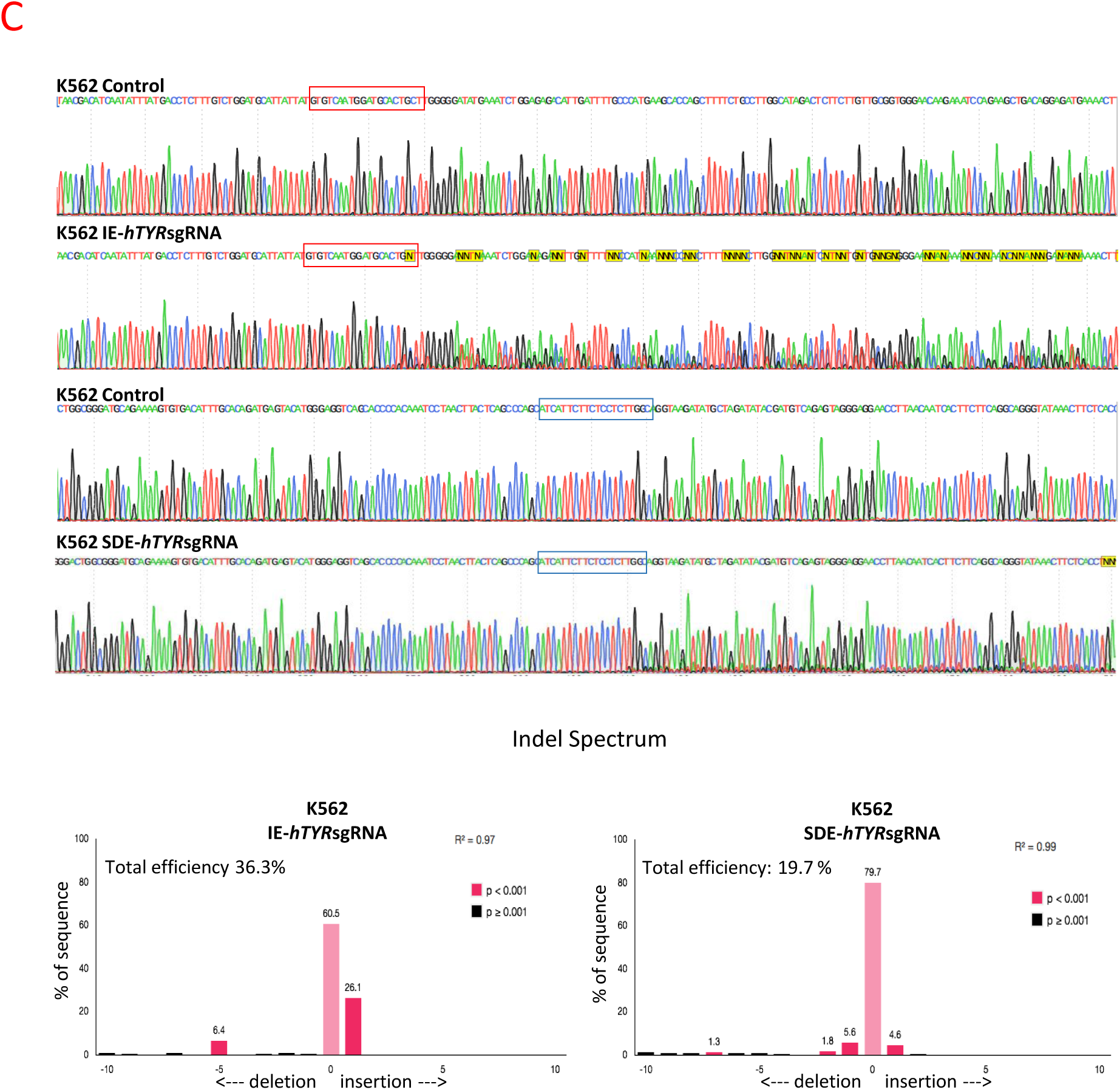
(C) CRISPR/Cas9 edited *TYR* sequences of K562 cells through IE-*hTYR*sgRNA (red box) and SDE-*hTYR*sgRNA (blue box). Control cells showed a wt sequence of the *TYR* gene, while K562-edited cells showed a mixture of sequences around the expected cleavage point for each sgRNA. TIDE analysis of sequences predicted the overall edition efficacy and most common allele variations generated.

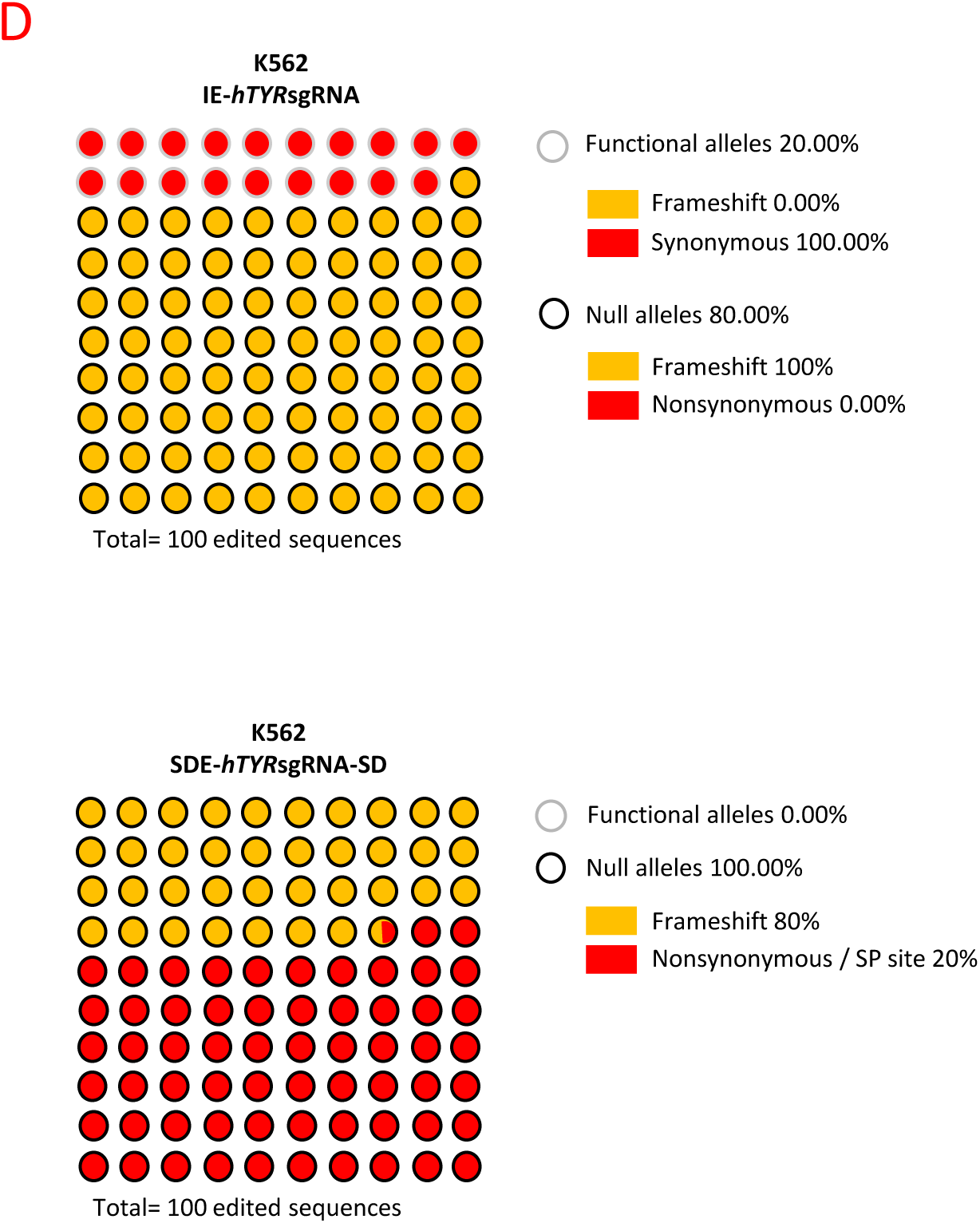
(D) NGS of *TYR* gene-edited K562 cells. Graphic representation of the mutations found, and their predicted effect on cells edited by IE-*hTYR*sgRN and SDE-*hTYR*sgRNA. Black and gray circles represent null and functional alleles, respectively, while the background indicates the type of mutation (red: nonsynonymous/sp site; yellow: frameshift). All SDE-*hTYR*sgRNA-edited sequences result in null alleles, in contrast with IE-*hTYR*sgRNA, which produced null alleles in only 80% of cases.

*In silico* analysis of mutated sequences obtained by NGS in IE-*hTYR*sgRNA K562-edited cells revealed that 20% of mutated alleles had nonsynonymous mutations that preserved the reading frame of the *TYR* gene (Figure 3D, Table 4). However, SDE-*hTYR*sgRNA was 100% efficient in generating null alleles, by inducing alterations in the sequence involved in splicing or frameshift-type mutations, or both (Figure 3D, Table 4).

**Table 4.**
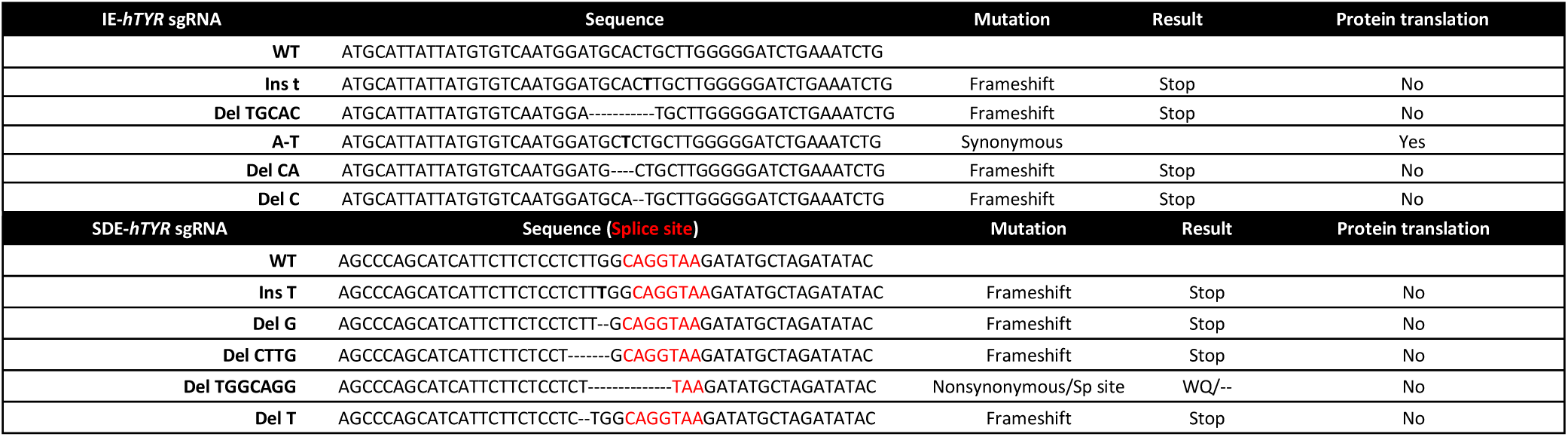
*In vitro* genomic edition of human *TYR* locus using sgRNA against exon coding sequence (IE) and coding SDE sequence. NGS analysis of allelic variants induced in K562 human cells.

### 3. sgRNA guides targeting the exon splice-donor site enhance knockout efficiency in other loci in human and mouse cells

To establish whether the knockout effect observed was Tyr-locus-dependent we decided to use the same strategy of targeting the ATM locus.

#### 3.1. *In silico* and NGS analyses of CRISPR/Cas9 sgRNA guides targeting the ATM exon splice-donor site identify a high frequency of null alleles in human and mouse cells

We investigated the effects of CRISPR-Cas9 DSBs induced in splice-donor sites of the ATM locus in human and mouse cells to address the putative universality of this process.

In a similar way as for the TYR locus, two custom-designed sgRNAs were used to modify the coding sequence of the *ATM* human gene in a central position of *ATM* exon 10 (IE-*hATM*sgRNA) and in the splice-donor sequence (SDE-*hATM*sgRNA) (Figure 4A). Three individual electroporation assays with each *ATM* sgRNA and empty vector were performed in K562 human cells. GFP expression was detectable 24 hours post-electroporation in all cases, indicating efficient CRISPR-Cas9 system delivery and expression (Figure 4B).

**Figure 4.**
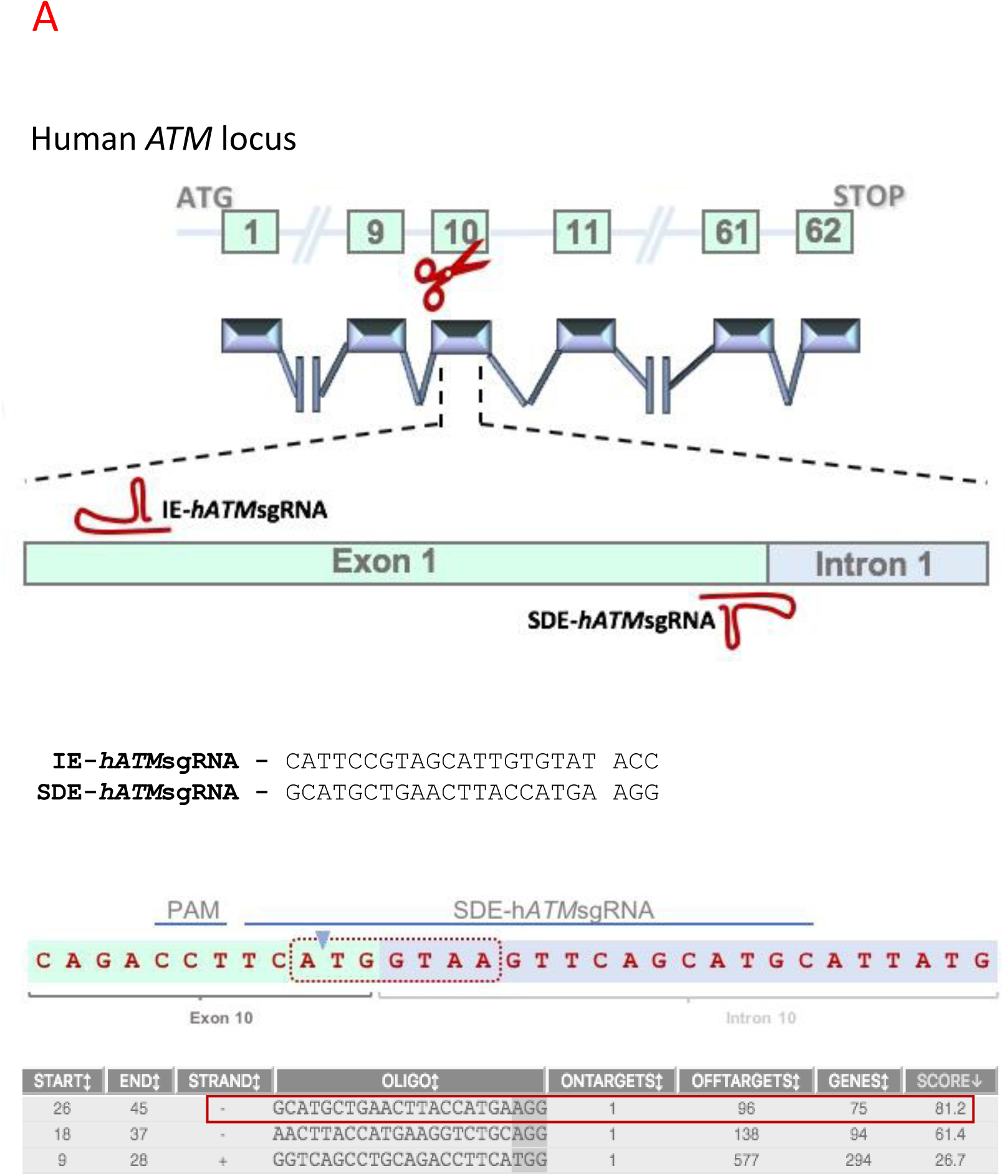
CRISPR/Cas9 editing of the human *ATM* locus. (A) Schematic representation of the human *ATM* locus and the two RNA guides represented in the exon 10 sequence. SDE-*hATM*sgRNA is complementary to the splice site between exon 10 and intron 10-11. IE-*hATM*sgRNA targeted a central position at the coding sequence of exon10. sgRNA sequences and their respective PAM sequences are also described. Exon-intron junction and nucleotides implicated in intron processing (red dot line). SDE-sgRNA (red box) was chosen among several candidates based on its score.

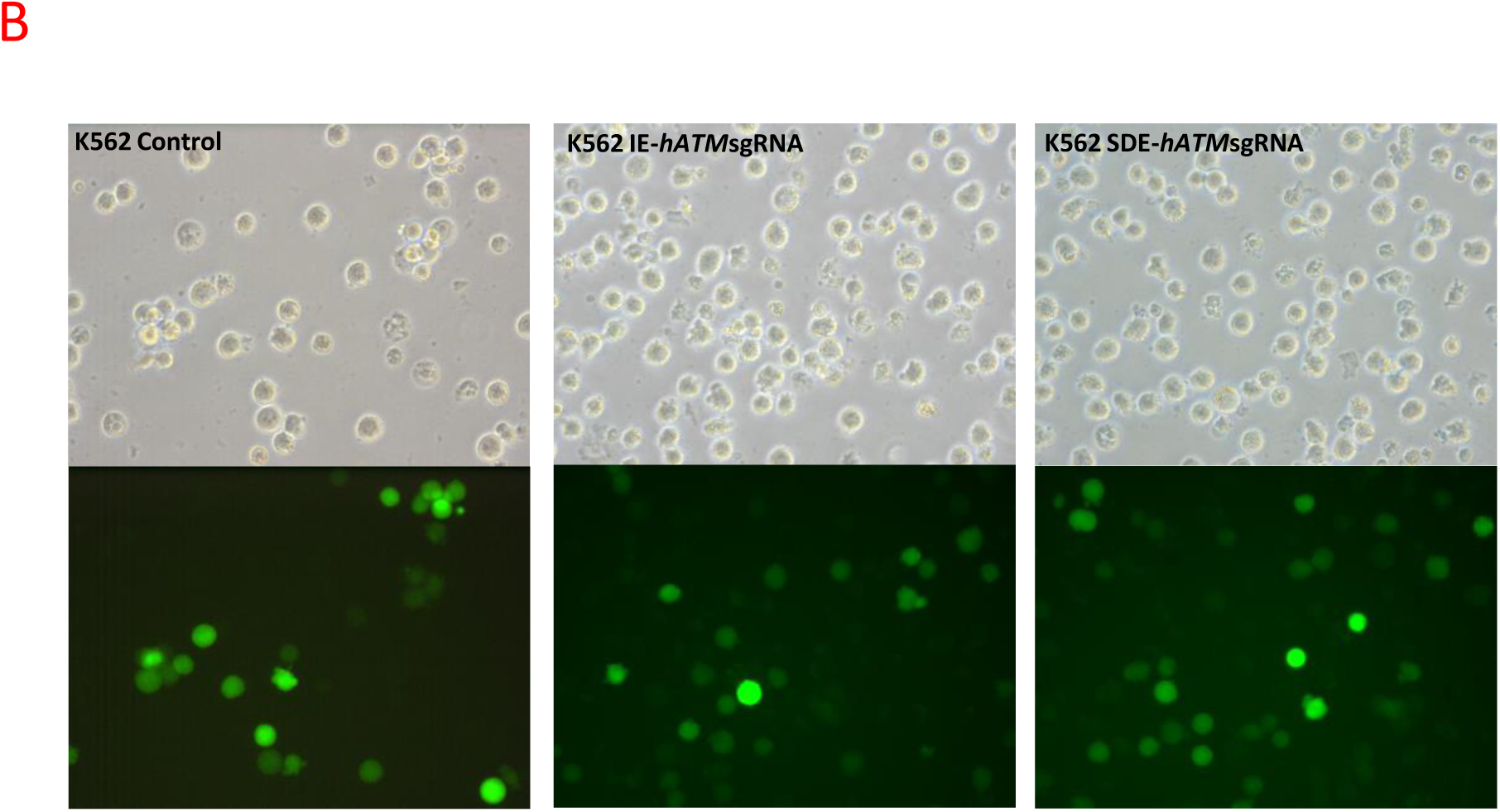
(B) CRISPR/Cas9 genome editing of the *ATM* gene in K562 human cells. Fluorescent microscopy of K562 human cells electroporated with empty px458 vector and carrying each RNA guides.

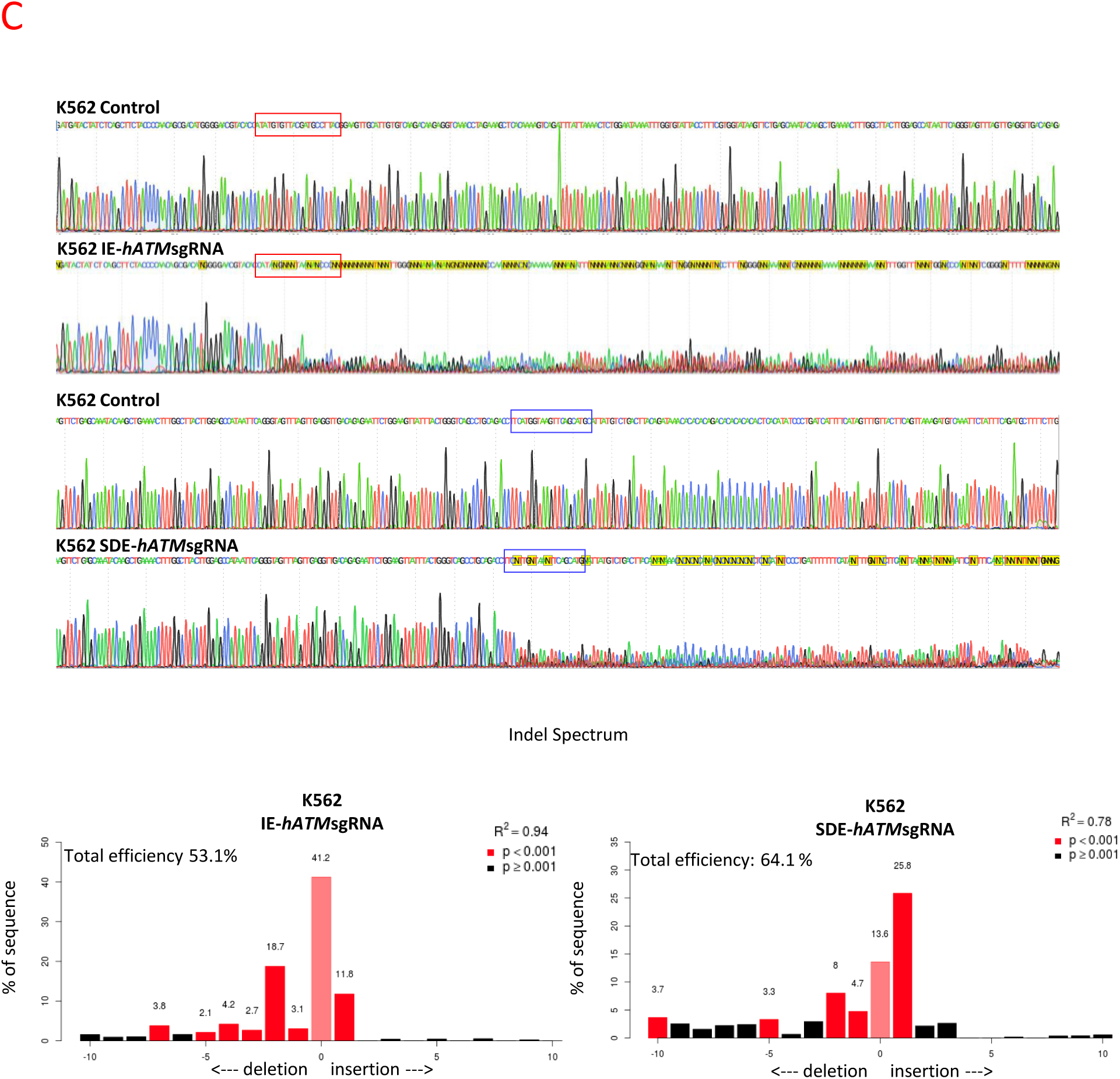
(C) CRISPR/Cas9-edited *ATM* sequences of K562 cells through IE-*hATM*sgRNA (red box) and SDE-*hATM*sgRNA (blue box). K562 cells expressing empty px458 vector, used as control, had a wild type sequence, while those expressing IE-*hATM*sgRNA (red box) and SDE-*hATM*sgRNA (blue box) showed a mixture of sequences around the expected Cas9 cleavage point. Lower panel illustrates the TIDE decomposition algorithm analysis of the edited sequences of *ATM* exon 10, showing highly efficient editing at the expected point.

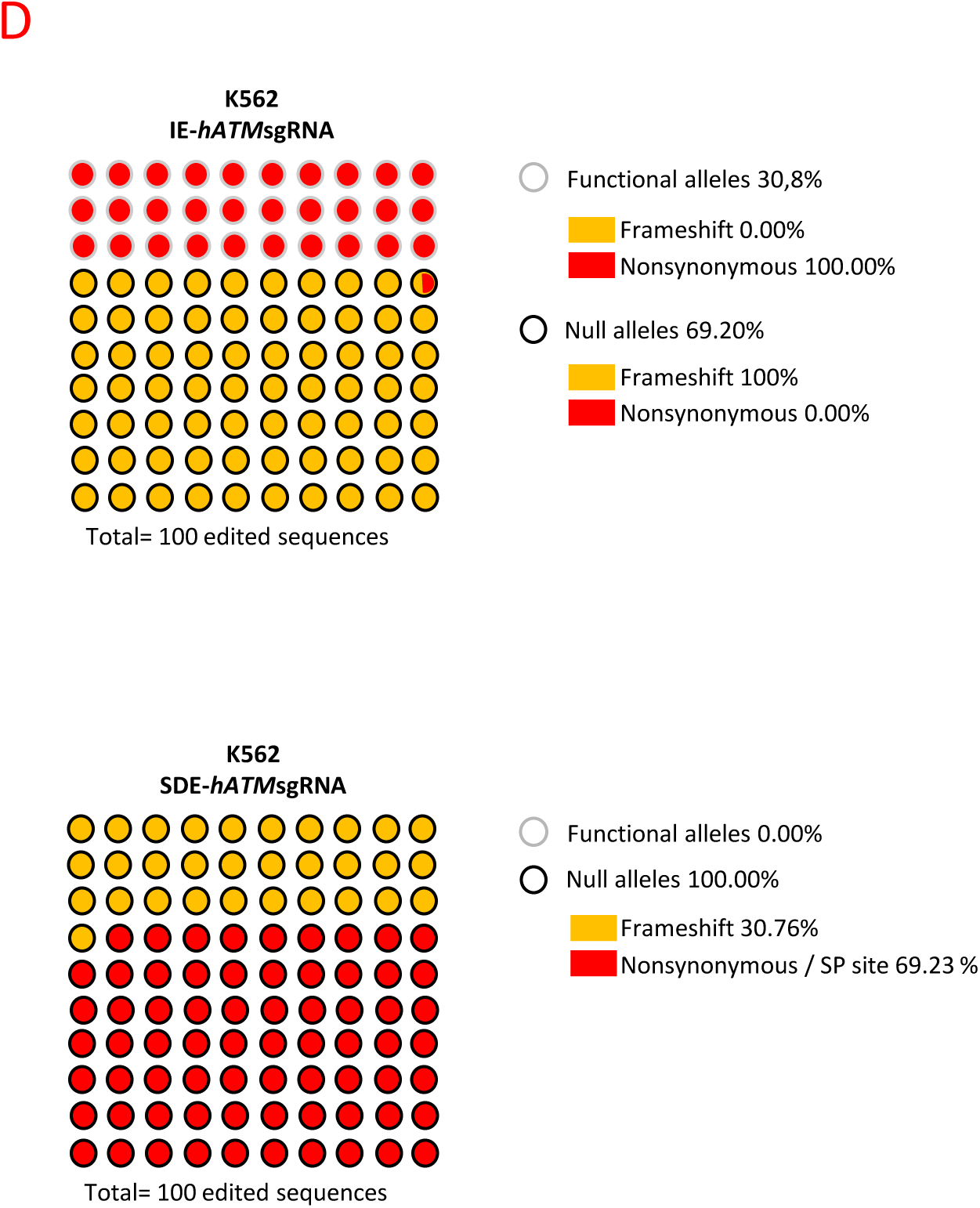
(D) NGS of edited K562 cells. Mutations found and their effect in edited cells predicted from IE-*hATM*sgRNA and SDE-*hATM*sgRNA. Black and gray circles correspond to null and functional alleles, respectively, while the background indicates the type of mutation (red: nonsynonymous/sp site; yellow: frameshift). The SDE-*hATM*sgRNA showed an increased efficiency in the generation of KO *ATM* alleles in K562 cells compared with IE-*hATM*sgRNA. Graphs indicate the percentage of edited sequences for each sgRNA, and which of them presumably give rise to null alleles. SDE-*hATM*sgRNA showed that 100% of edited sequences would generate null alleles, while only 69.20% of IE-*h*ATMsgRNA-edited sequences produce null alleles.

Sanger sequencing identified indel mutations at the expected cleavage point in knockout assays, while no changes in sequence were observed in control cells (Figure 4C). TIDE analysis showed similar overall editing efficiency for IE-*hATM*sgRNA (53.10%) and SDE-*hATM*sgRNA (64.10%) in the K562 cell line. The TIDE algorithm predicted different indels without the predominance of any lengths in IE-*hATM*sgRNA and SDE-*hATM*sgRNA-edited cells (Figure 4C).

The mutated ATM sequences generated in IE-*hATM*sgRNA and SDE-*hATM*sgRNA-edited K562 cells were analyzed by NGS (Table 5). The mutations detected and their predicted effect in both cases corroborated the TIDE prediction. Excluding wildtype alleles, *in silico* analysis of mutant alleles in SDE-*hATM*sgRNA predicted a null effect in all of them. Nevertheless, null mutations (69.2%) and nonsynonymous mutations preserving the reading frame (30.8%) were detected with IE-*hATM*sgRNA. All null mutations detected with IE-*hATM*sgRNAs were produced by frameshift mutations (Figure 4D, Table 5).

**Table 5.**
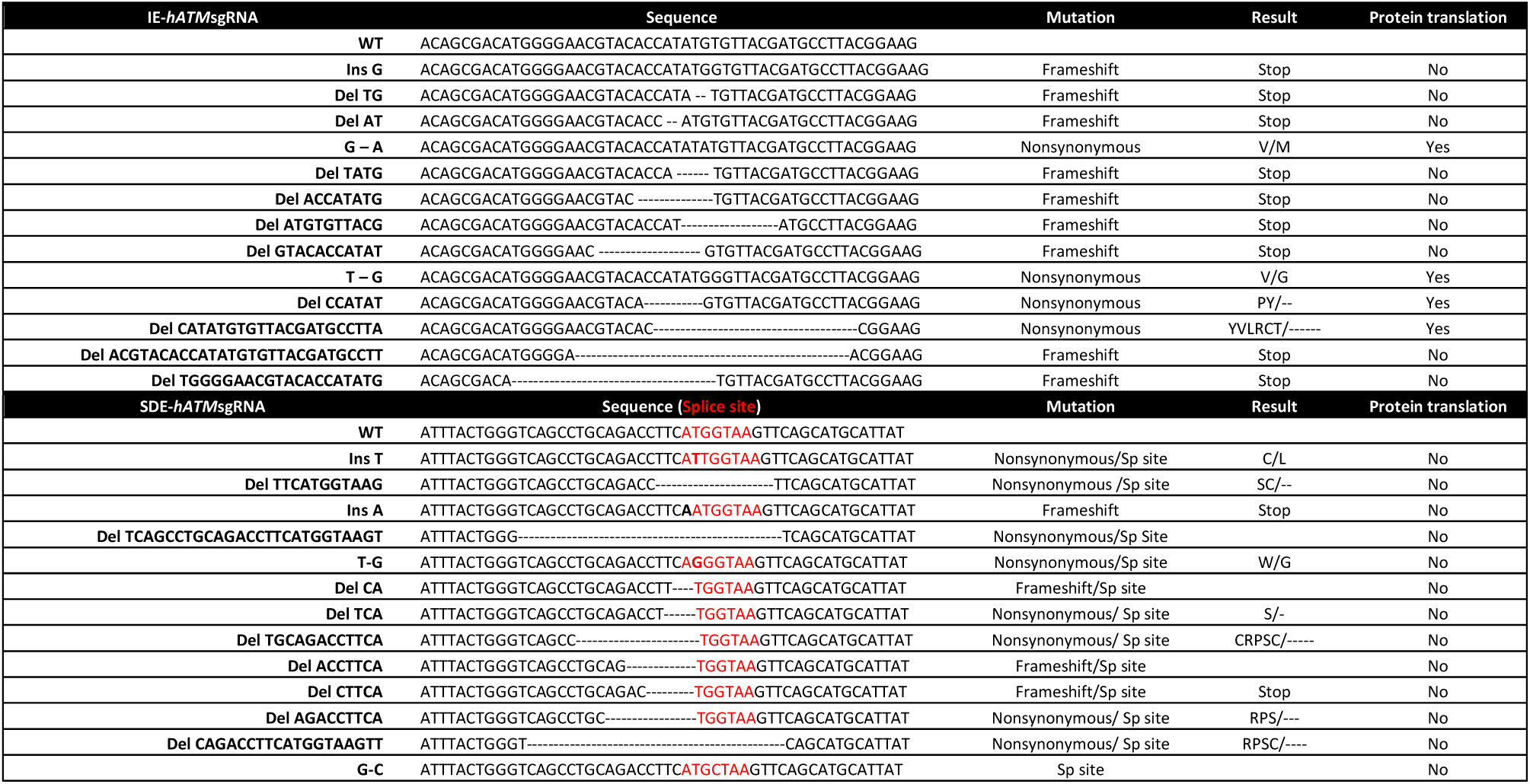
*In vitro* genome editing of the human *ATM* locus using sgRNA against the exon coding sequence (IE) and the coding SDE sequence. NGS analysis of allelic variants induced in K562 human cells.

To study whether the effects observed in K562 cells were specific to human, an analog design was applied to exon 10 of the mouse *Atm* gene. Similarly, two sgRNA guides were designed in exon 10 (IE-*mAtm*sgRNA) and in the coding sequence involved in the intron splicing processing (SDE-*mAtm*sgRNA) (Figure 5A). Three individual electroporation assays with each *ATM* sgRNA and empty vector were performed in Baf/3 mouse cells. GFP expression was detectable 24 hours post-electroporation in all cases, indicating an efficient CRISPR-Cas9 system of delivery and expression (Figure 5B). Sanger sequencing revealed the presence of indel mutations at the expected cleavage point in knock-out assays, while no sequence changes were observed in control cells (Figure 5C). The TIDE algorithm predicted small indels (1-7 bp) in IE-*mAtm*sgRNA and SDE-*mAtm*sgRNA-edited sequences (Figure 5C).

**Figure 5.**
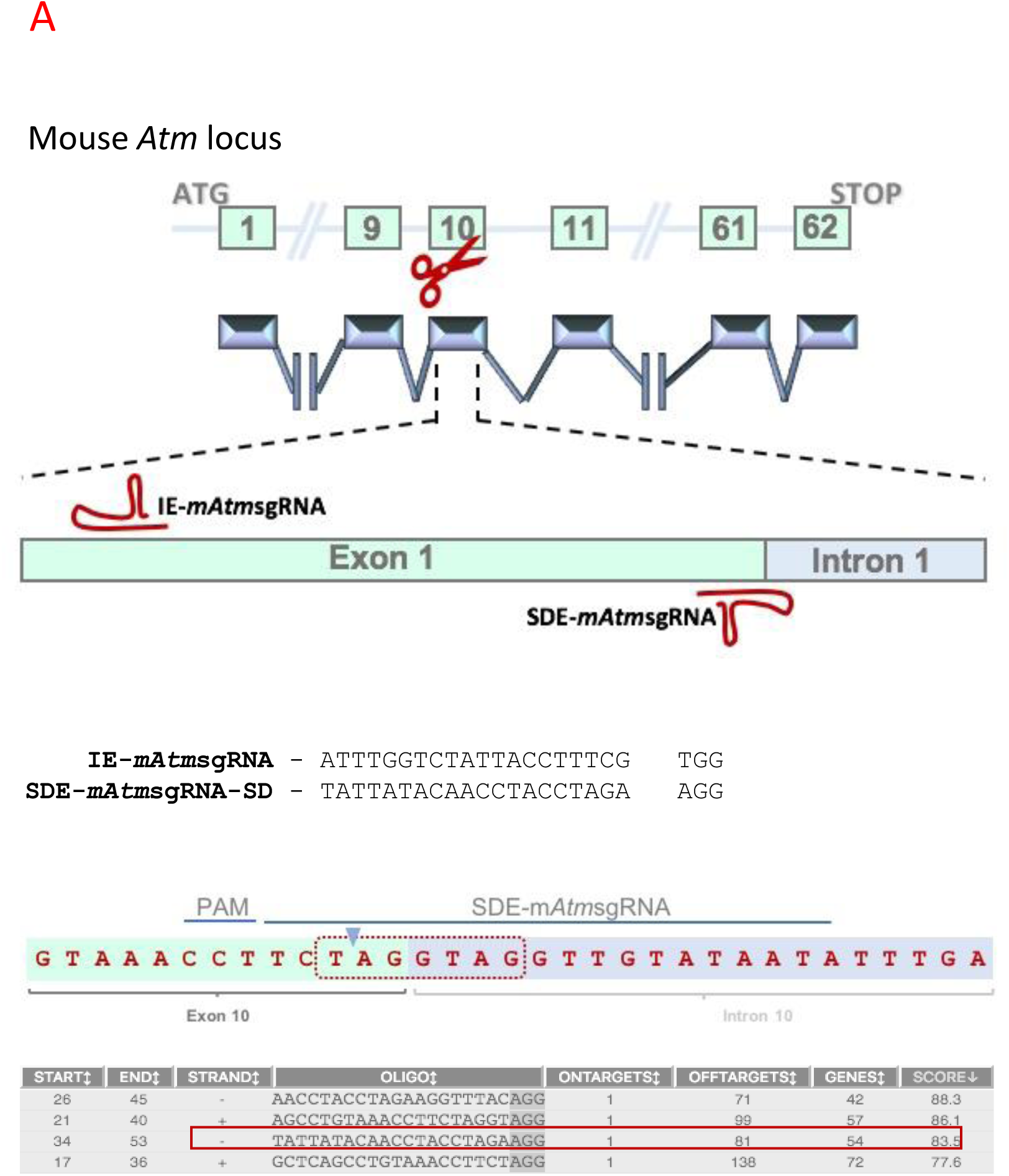
CRISPR/Cas9 editing of the mouse *Atm* locus. (A) Schematic representation of the mouse *Atm* locus and the two RNA guides represented in the exon 10 sequence. SDE-*mAtm*sgRNA is complementary to the splice site between exon 10 and intron 10-11. IE-m*Atm*sgRNA is located in the coding sequence of exon10. sgRNA sequences and their respective PAM sequences are also described. Exon-intron junction and nucleotides implicated in intron processing (red dot line). SDE-sgRNA (red box) was chosen among several candidates based on its score.

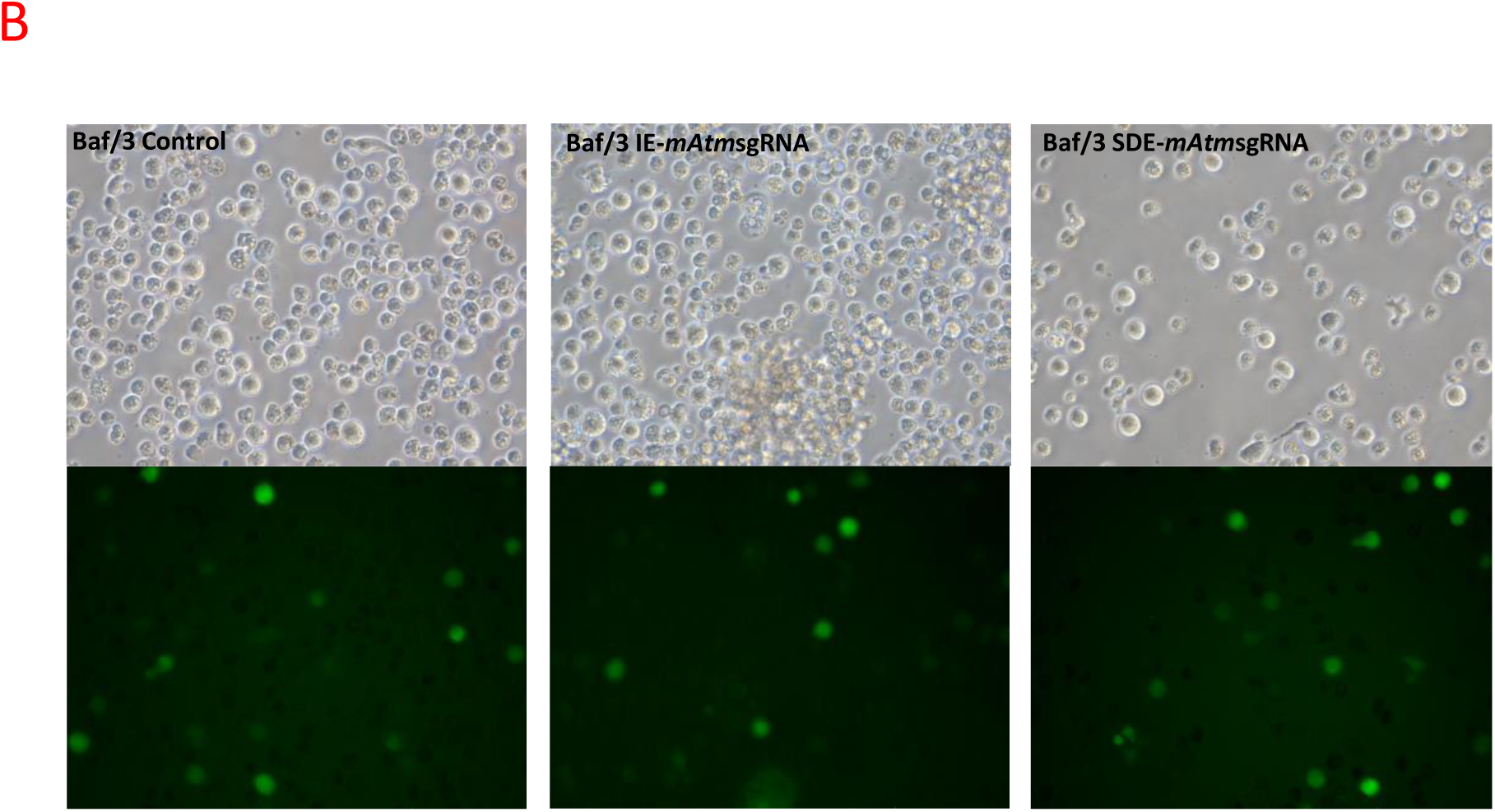
(B) CRISPR/Cas9 genome editing of the *Atm* gene in Baf/3Baf/3 mouse cells. Fluorescent microscopy of Baf/3Baf/3mouse cells electroporated with empty px458 vector and carrying each RNA guides.

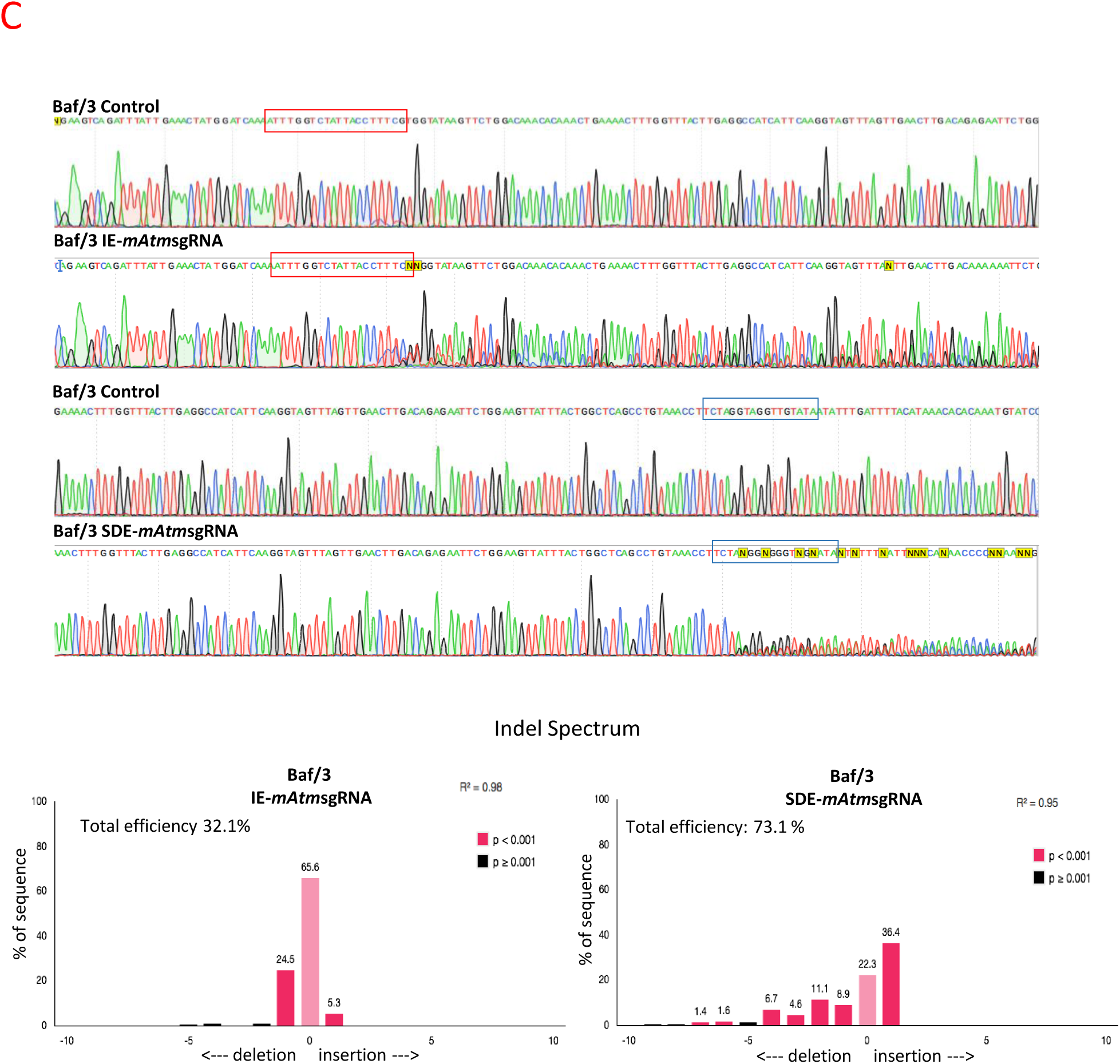
(C) CRISPR/Cas9-edited *Atm* sequences of Baf/3Baf/3 cells through m*Atm*sgRNA-IE (red box) and SDE-*mAtm*sgRNA (blue box). Baf/3cells expressing empty px458 vector, used as control, had a wild type sequence, while cells expressing IE-*mAtm*sgRNA (red box) and SDE-*mAtm*sgRNA (blue box) showed a mixture of sequences around the expected Cas9 cleavage point. Lower panel shows TIDE decomposition algorithm analysis of the edited sequences of *Atm* exon 10, indicating highly efficient editing at the expected point

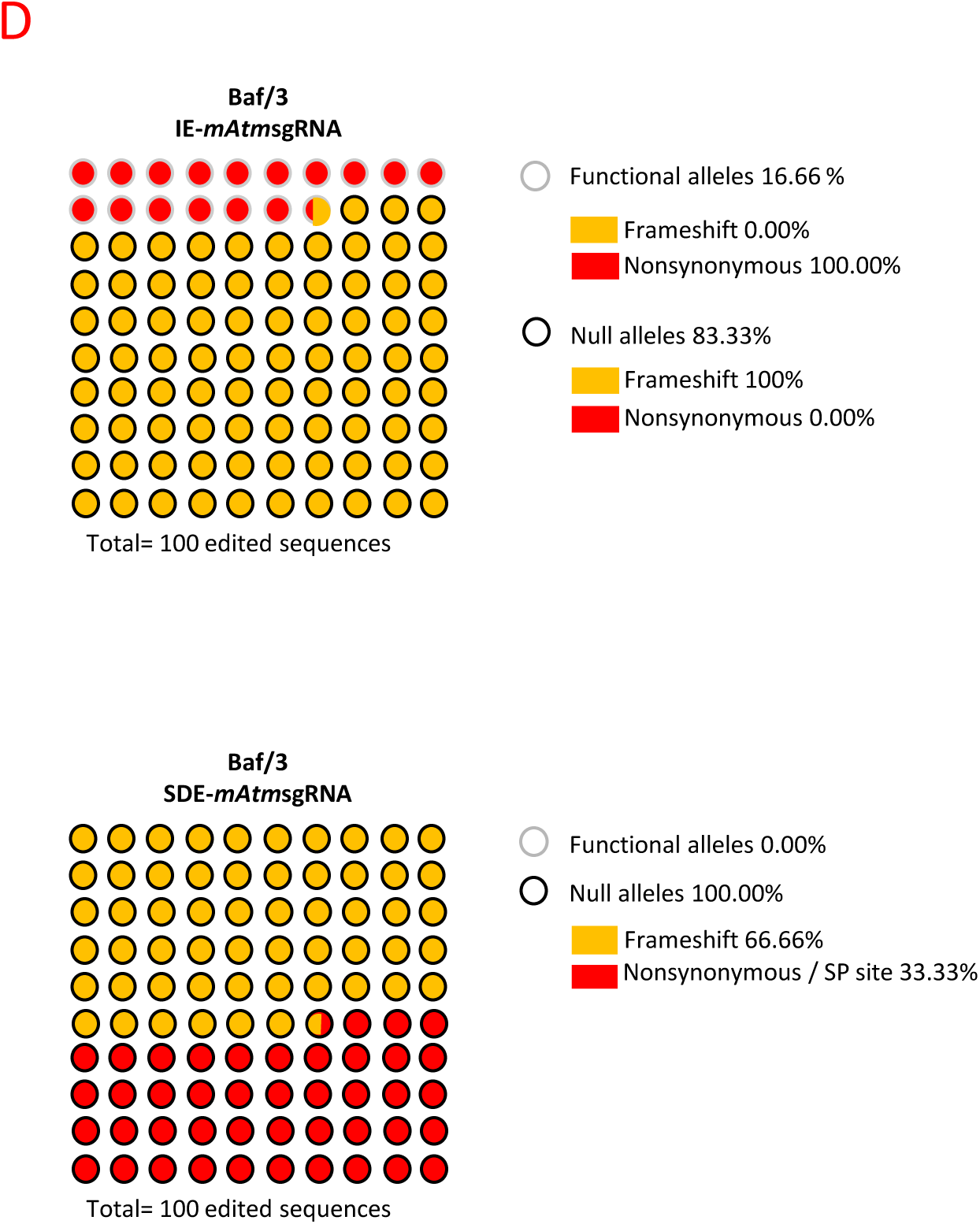
(D) NGS of edited Baf/3 cells. Mutations identified and their predicted effect in edited cells as revealed by IE-*mAtm*sgRNA and SDE-*mAtm*sgRNA. Black and gray circles correspond to null and functional alleles, respectively, while the background indicates the type of mutation (red: nonsynonymous/sp site; yellow: frameshift). The SDE-*mAtm*sgRNA was more efficient at generating knockout *Atm* alleles in Baf/3 cells compared with IE-*mAtm*sgRNA-IE. Graphs illustrate the percentage of edited sequences arising from each sgRNA, and which of them presumably give rise to null alleles. SDE-*mAtm*sgRNA showed that 100% of edited sequences would generate null alleles, while IE-*mAtm*sgRNA-IE indicate that only 83.33% of edited sequences would produce null alleles.

NGS analysis of the IE-*mAtm*sgRNA-edited mouse *Atm* locus revealed six generated mutated alleles, 83.33% of which resulted in null alleles, due to frameshift mutation, and 16.67% of which maintained their functionality because they harbored a nonsynonymous mutation. In contrast, SDE-*mAtm*sgRNA was able to induce frameshift and/or nonsynonymous mutations in all cases (Figure 5D, Table 6), but gave rise to null alleles by altering the canonical splice sequence.

**Table 6.**
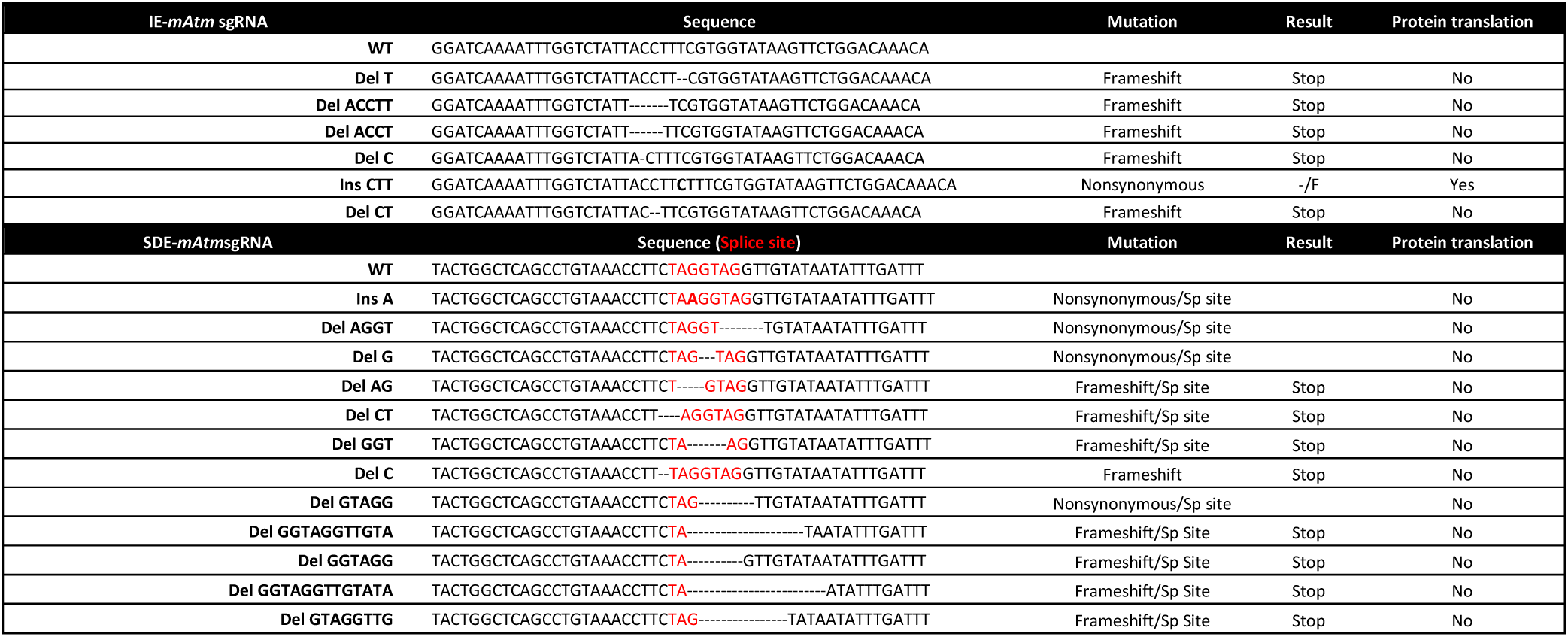
*In vitro* genome editing of the mouse *Atm* locus using sgRNA against the exon coding sequence (IE) and the coding SDE sequence. NGS analysis of allelic variants induced in Baf/3 mouse cells.

#### 3.2 Functional analysis

ATM protein levels in K562-edited cells were analyzed by western blot (WB) to evaluate the functionality of the mutant alleles generated by the CRISPR/Cas9 system in the human ATM gene (Figure 6A). While IE-*hATM*sgRNA-transfected cells showed slightly weaker ATM expression compared with K562 parental cells, low levels of ATM protein were detected in SDE-*hATM*sgRNA-transfected cells (Figure 6A). Single-cell-derived cell lines from both IE-*hATM*sgRNA (6 clones) and SDE-*hATM*sgRNA-SD (6 clones) K562 cells were established and analyzed by NGS (Table 7). ATM protein levels of each single-cell-derived clone were analyzed by WB. Most mutated cell clones (4/6) edited with IE-*hATM*sgRNA showed ATM expression (Figure 6B). NGS analysis of all single-cell clones edited with IE-*hATM*sgRNA had at least one functional allele, either a wildtype (wt) or with nonsynonymous mutation (Figure 6B, Table 7). However, several mutated cell clones (5/6) edited with SDE-*hATM*sgRNA had no levels of ATM protein that could be detected by WB (Figure 6B). Analyzing them showed splicing mutations together with nonsynonymous or synonymous mutations in both *ATM* alleles (Table 7).

**Figure 6.**
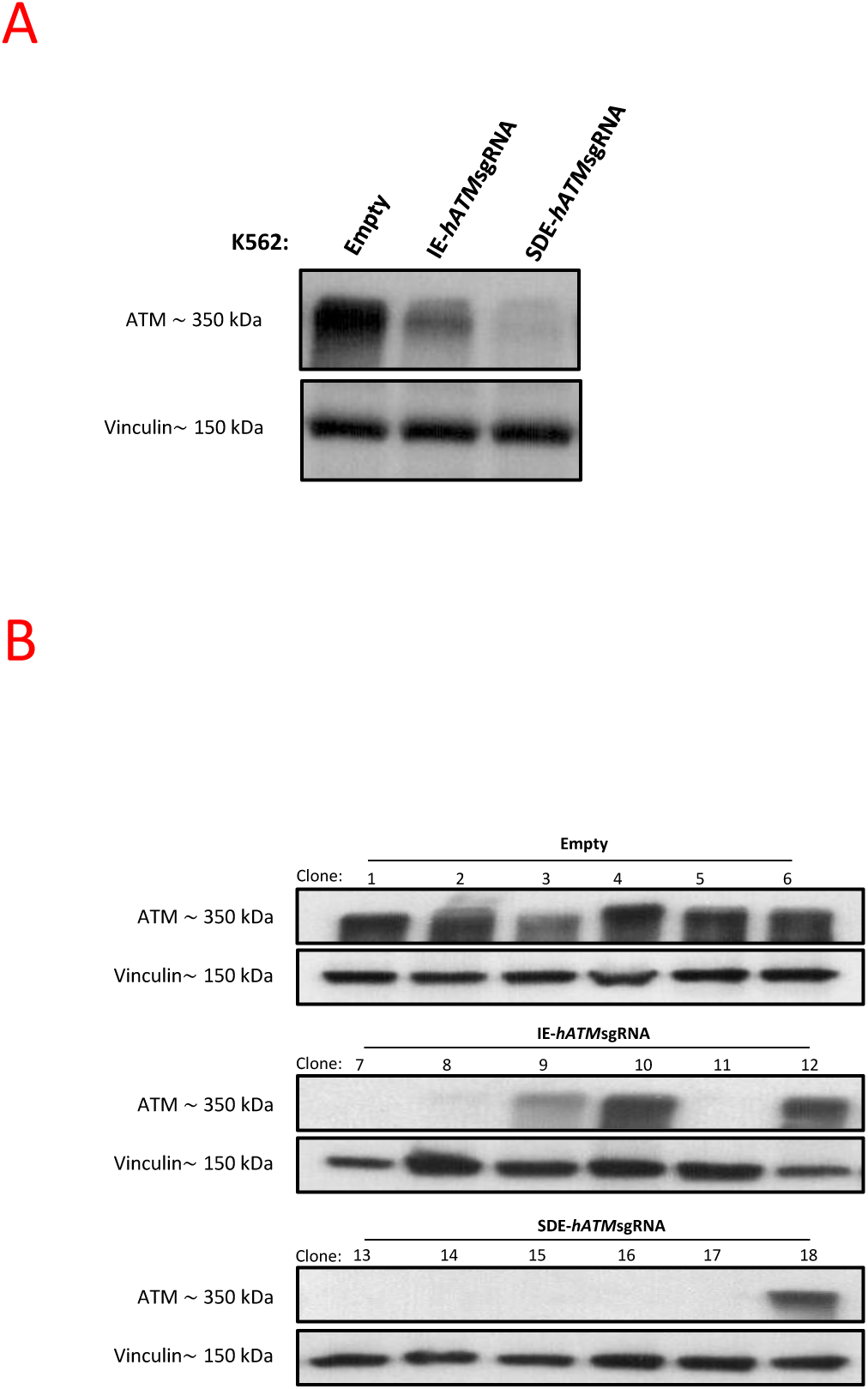
Analysis of *ATM* null-allele generation by CRISPR/Cas9-induced mutations in K562 cells. (A) Western blot analysis of *ATM* protein expression in K562-edited cells. A single band of 350 kDa corresponding to ATM was observed in K562 cells electroporated with empty px458. A lower level of ATM expression was observed in IE-*hATM*sgRNA-edited cells, and an even lower level was noted in SDE-*hATM*sgRNA-edited cells. Vinculin expression of the cells was used as the loading control. (B) Western blot analysis of ATM expression in single-edited-cell clones. All clones derived from cells electroporated with empty vector, used as a control, showed a single band corresponding to ATM. Three of six IE-*hATM*sgRNA edited clones showed no expression of ATM and one of six had a lower level of ATM expression compared with controls. Only one of six SDE-*hATM*sgRNA-edited clones expressed ATM, while ATM expression could not be detected in the other five clones.

**Table 7.**
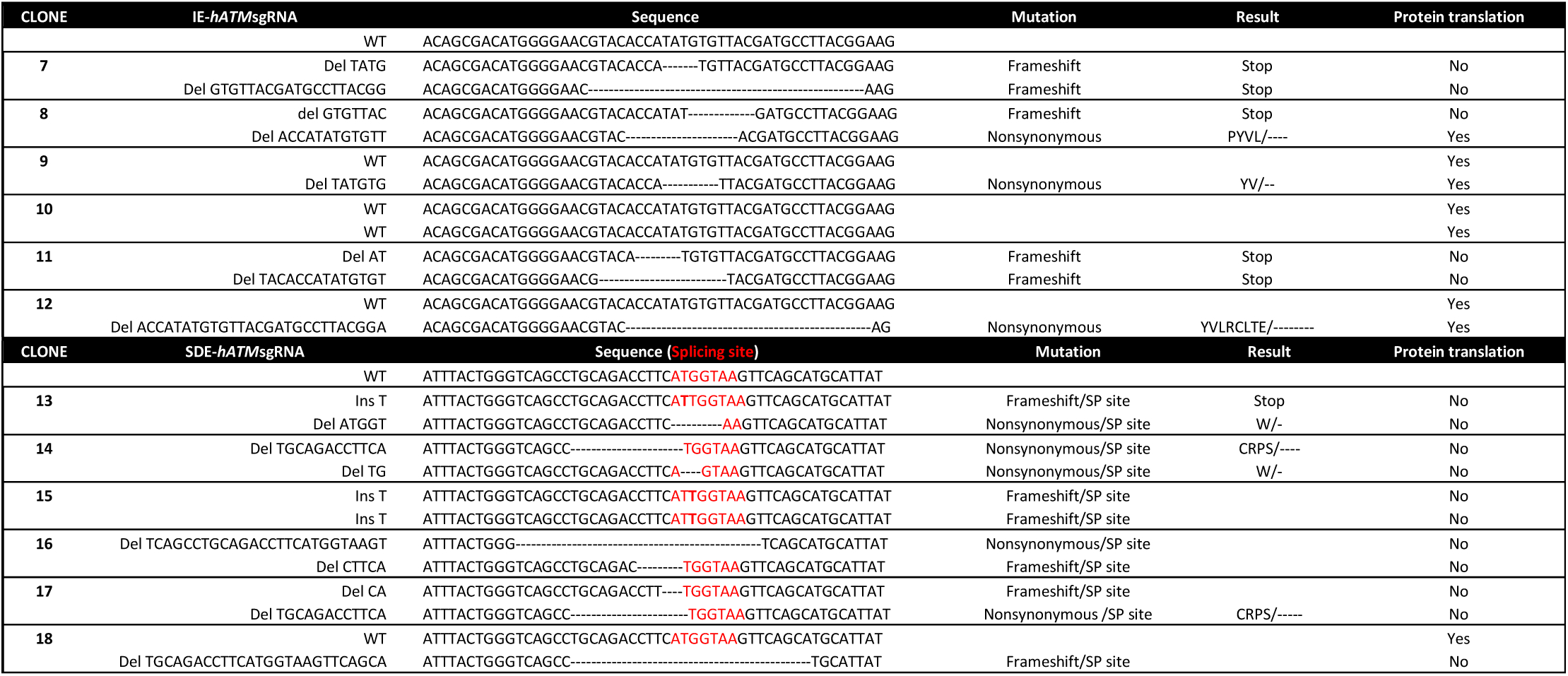
NGS analysis of *ATM* allelic variants induced in human K562 single-edited cell-derived clones.

### 4. The sgRNA guide targeting the exon splice-donor site of ABL increases the yield of null alleles and enhances BCR/ABL oncogene knockout efficiency in the K562 BCR/ABL cell line

Various strategies at different molecular levels(27) have been employed to treat malignant diseases in recent decades, such as specific drug inhibitors acting at the protein level, gene suppression therapies at the mRNA level, and genome-editing nucleases at the DNA level. CRISPR/Cas9 works at the DNA level and has the advantage of providing permanent and full gene knockout, while mRNA or protein-inhibiting approaches only work at the mRNA level. To test the efficiency of SDE-sgRNA and IE-sgRNA guides at switching off oncogenes we performed similar assays to generate ABL null alleles in the leukemic K562 cell lines and to abrogate the oncogene activity of BCR/ABL oncogene fusion.

### 4.1. The sgRNA guides targeting the exon 1 splice-donor site of ABL enhance the generation of null ABL alleles

Three individual electroporation assays of K562 cells were performed with each sgRNA directed towards the ABL exon 1 (SDE-*hABL-1*sgRNA and IE-*hABL-1*sgRNA) cloned in a CRISPR-Cas9-GFP mammalian expression vector. An empty CRISPR-Cas9-GFP vector was used as a control. GFP expression was detectable 24 hours post-electroporation in all cases, indicating effective delivery of the CRISPR/Cas9 system and its expression in K562 cells (Data not show). Sanger sequencing of K562 showed genome edition at expected cleavage point for each sgRNA guide and Tide analysis predicted a variety of small indels for each guide (Figure 7A).

**Figure 7.**
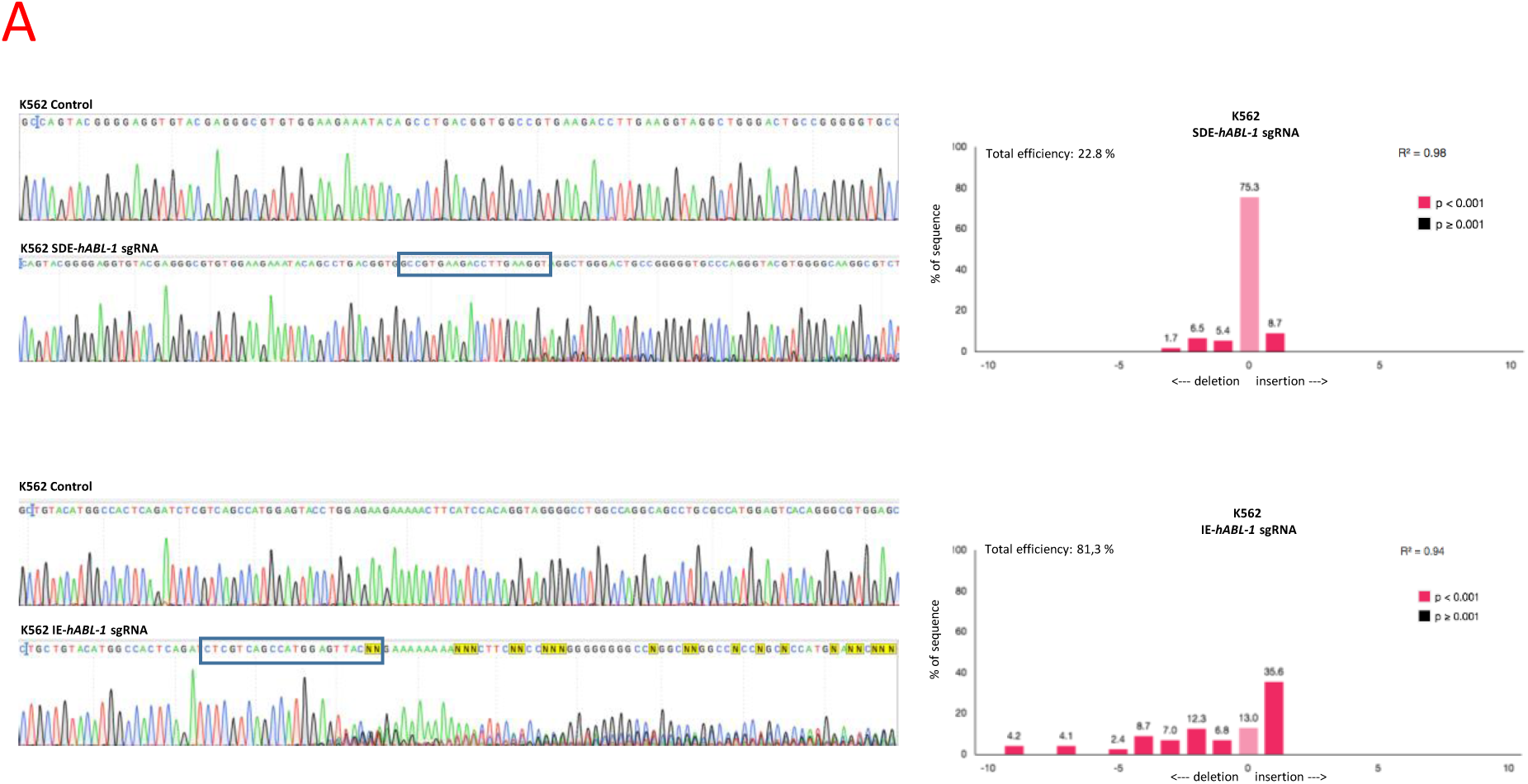
Functional analysis of BCR-ABL-1 null-allele generation by CRISPR/Cas9-induced mutations in K562 cells. (A) Analysis of SDE and IE-*hABL*-1sgRNAs genomic editing in K562 cells. Sanger sequencing of ABL-1 exon 6 showed a mixture of sequences at the expected cleavage point in both sgRNAs. TIDE algorithm analysis predicted more frequent mutations for each sgRNA.

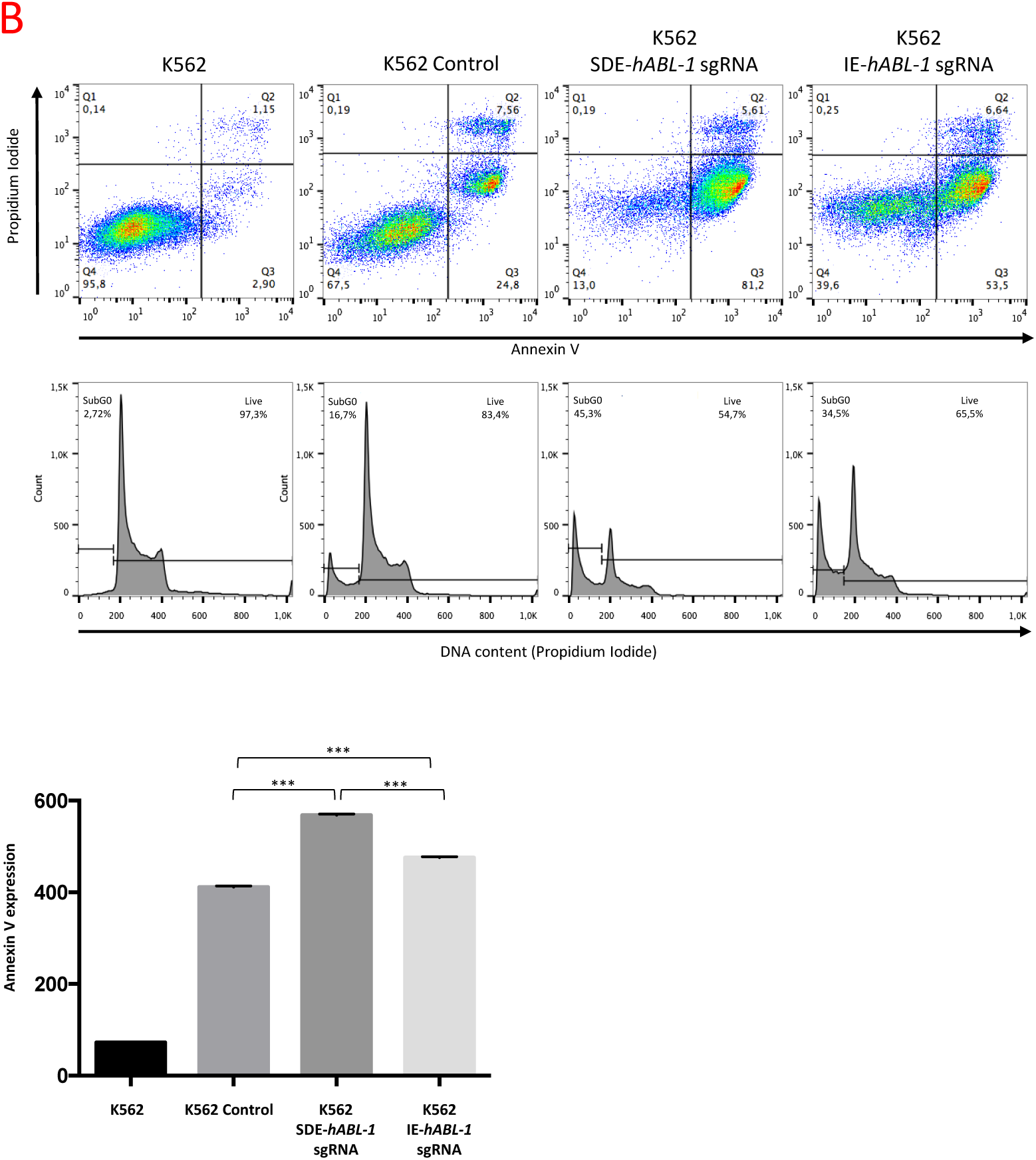
(B) Flow cytometry analysis of annexin V expression and cell cycle of K562-edited cells. SDE-*hABL-1*sgRNA-edited cells had a higher level of apoptosis than K562 cells edited by IE-*hABL-1*sgRNA and control cells after electroporation with the empty vector. The DNA content of the cells edited with SDE sgRNA gave 10% higher levels than IE-edited cells (45.3% *vs.* 34.5%). Plots show results of a representative experiment from three independent replicates. The quantification of annexin expression in K562-edited cells with SDE and IE hABL-1 sgRNAS showed a higher level of expression in SDE-hABL-1sgRNA edited cells (568,2 mfi) than in IE-hABL-1sgRNA-edited cells (475.5 mfi). Graph shows results from three independent experiments. ***, p<0.001.

NGS analysis showed the most frequent allele variations generated in K562 by electroporation with SDE-and IE-*hABL*-1 sgRNAs (Table 8). 40% (4/10) of the allelic variations generated by IE-*hABL*-1 sgRNA gave rise to nonsynonymous mutations. By contrast, SDE-hABL-1 sgRNA gave rise to 100% (9/9) of knockout sequences (Table 11), four of which (44.4%) were nonsynonymous mutations, but with an altered canonical splicing sequence (Table 8).

**Table 8.**
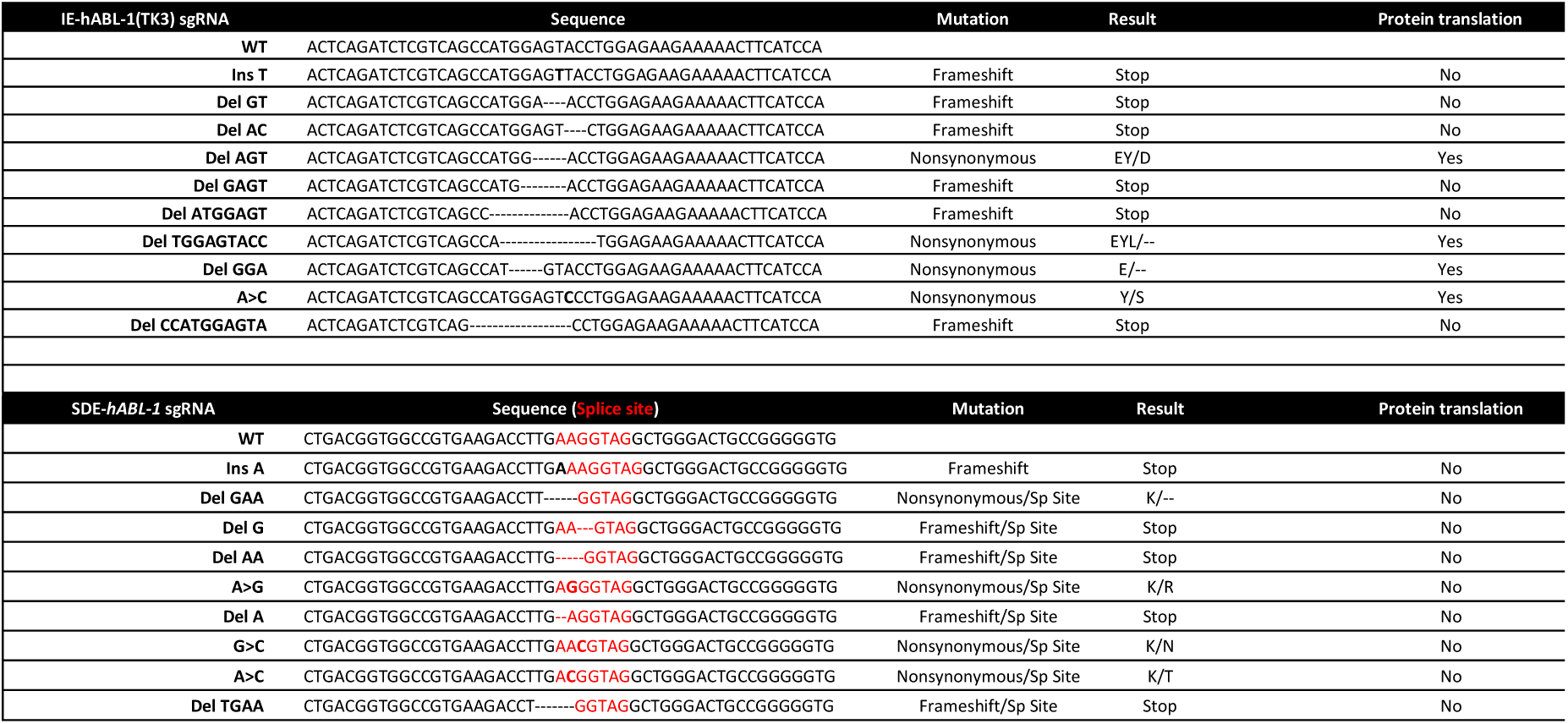
*In vitro* genome editing of the human ABL-1 locus using sgRNA against the exon coding sequence (IE) and the coding SDE sequence. NGS analysis of allelic variants induced in K562 cells.

### 4.2. The SDE-*hABL-1*sgRNA enhances the efficiency for abrogating cell survival/ proliferation BCR-ABL dependent

To test the ability of SDE sgRNAs to increase the efficiency at knocking out fusion oncogenes, we compared the proficiency at abrogating the cell survival and proliferation produced by the BCR-ABL oncoprotein through the induction of indels with SDE-sgRNA and IE-sgRNA CRISPR-Cas9 guides.

In three independent assays, we electroporated the K562 BCR/ABL cell line with SDE-hABL-1 and IE-hABL-1sgRNA. After 48 hours, we analyzed the effects on apoptosis and cell cycle (Figure 7B). SDE-*hABL-1*sgRNA-targeted cells showed a higher level of apoptosis (86.8%) than noted in IE-*hABL-1*sgRNA cells (60.1%), while 32.4% of control cells were apoptotic. K562 cells electroporated with SDE sgRNA yielded 10% more subG0 DNA content (45.3%) than IE-edited cells (34.5%) (Figure 7B). The quantification of annexin expression in K562-edited cells with SDE-and IE-*hABL-1* sgRNAS showed a higher level of expression in SDE-*hABL*-1sgRNA-edited cells (568.2 mfi) compared with IE-hABL-1sgRNA-edited cells (475.5 mfi) and K562 control cells (411.5 mfi) (Figure 7B).

## DISCUSSION

DSB induced by CRISPR/Cas9 technology is the gold standard for creating null alleles in any biological system. In most cases, DSBs are typically repaired by non-homologous end-joining (NHEJ), resulting in indel mutations. These mutations can generate knockout alleles when CRISPR/Cas9 is directed at coding sequences, but due to the variable size of NHEJ-induced indels, generating a full KO in one step cannot always be achieved at high frequency. In fact, full KO generation requires off-frame mutations in both alleles and this is a random question because several mutations could preserve the reading frame (e.g., +3 or −3 mutations). This may be a minor matter if the target cell can be selected, sequenced and grown. However, in cancer gene therapies relying on the disruption of a driver oncogene, a delivery strategy resulting in the expression of both Cas9 nuclease and the knockout sgRNA in transduced/transfected cancer cells is desirable. This is especially critical for *in vivo* approaches and for *in vitro* gene therapy where the expansion processes from a selected edited cell are not available. If there is an acceptable degree of efficiency of delivery of CRISPR/Cas9 reagents to the cancer cell, the key step for success lies in the effectiveness of a specific sgRNA at knocking out the oncogene. In this cellular context, the null effect could be increased by sgRNAs targeting the exon SD boundaries. Following this strategy, the generation of null alleles could be increased in two independent ways: by the probabilities of producing a frameshift mutation and/or breaking the canonical pre-mRNA splicing. In the present work we have demonstrated that knockout efficiency can be increased using sgRNAs targeting the exon splice donor area. The study considered the predicted informatic score (most guides with a score of > 80) and the cut-site of the sgRNAs. It is important to note that for SDE-sgRNAs we chose PAMs to trigger DSBs inside the coding sequence that were located no further than four nucleotides from the end of the exon.

We noted that most of the mutant alleles produced in our assays in the Baf3 and k562 cell lines correspond to small indels, indicating that the DSB is repaired by blunt-end ligation independently of sequence homology, the classic nonhomologous end joining (C-NHEJ) mechanism(7). NGS corroborated the Sanger sequences detected and exposed new mutant alleles that are likely to be little-represented in the edited cell line. As expected, NGS and Sanger sequencing highlighted the same alleles in *in vivo* assays of mouse zygotes, grown to blast or of mice born from them. *In silico* analysis of these mutant alleles revealed a full efficiency of the null effect in SDE-sgRNA compared with IE-sgRNA. When an IE-sgRNA was used, mutant alleles with mutations preserving the reading frame were detected. To corroborate the *in silico* findings we Sanger-sequenced all mice born in both groups. Excluding unmutated mice, we detected color mice born from microinjected zygotes with IE-sgRNA with indels in one or more alleles. It is of particular note that we observed color mice with both alleles mutated, one of them with a frameshift mutation and the other with a mutation, indicating that some induced indels are not able to generate a frameshift mutation. By contrast, when we used a Tyr SDE-sgRNA, we detected albino or mosaic mice featuring one allele with a frameshift mutation and another with a mutation but a destroyed splice-donor site. This result demonstrates the higher null efficiency when an SDE-sgRNA is used. To determine whether this effect can be reproduced in another locus we employed the same assay but targeting the *ATM* and *ABL* loci. A similar result was obtained in both loci in human and mouse cell lines. Western blot analysis in cell clones from both groups corroborated the NGS and the results of their *in silico* analysis. More importantly, this approach can be efficiently used to abrogate oncogene expression. When a cancer cell is the target, a delivery strategy that can result in the expression of Cas9 and an oncogene-specific sgRNA in all infected cells is desirable. This is especially critical for *in vitro* gene therapy where the expansion processes of a selected edited cell are not available. Similarly, it is crucial for *in vivo* approaches in cancer therapies based on disrupting a driver oncogene. If the efficiency of delivery of CRISPR/Cas9 reagents to the cancer cell is acceptable, the key step for success lies in the effectiveness of a specific sgRNA at knocking out the oncogene. In most of these cases, the designs are based solely on off-target criteria. However, for those cases in which cellular selection is not an option and only one sgRNA can be used, the null effect could be increased with an sgRNA targeting the exon boundary. Following this strategy, we abrogated p210 (BCR/ABLp210) oncoprotein expression in the K562 cell line. Using this approach, pools of K562 edited cells electroporated with SDE-sgRNAs or IE-sgRNA were studied. The loss of p210 expression in K562 cells with SDE-sgRNA resulted in a significant increase in apoptosis levels. Thus, this strategy could be adopted for gene therapy in cases for which cell selection is not an option and the delivery Cas9 vector only allows the accommodation of one sgRNA.

## CONCLUSIONS

Genome-editing nucleases, like the popular CRISPR/Cas9, enable knockout cell lines and null zygotes to be generated by inducing site-specific DSBs within a genome. In most cases, when a DNA template is not present, the DSB is repaired by non-homologous end joining, resulting in small nucleotide insertions or deletions that can be used to construct knockout alleles. However, for several reasons, these mutations do not produce the desired null result in all cases, giving rise to a similar but functionally active protein. This undesirable effect could limit the efficiency of gene therapy strategies based on abrogating oncogene expression by CRISPR/Cas9 and should therefore be borne in mind. The use of an sgRNA-targeting splice donor site could improve the null result for *in vivo* gene therapies. This strategy could be adopted to abrogate *in vivo* the oncogenic activity involved in tumor maintenance.

## ACKNOWLEDGMENTS

The authors wish to express their sincere thanks to Dionisio Martín, Alberto Pendás (Spanish Research Council, CSIC), for their technical assistance and contribution to our CRISPR/Cas9 studies; Servicio de Citometría, Servicio de Experimentación Animal (University of Salamanca) and Servicio de Secuenciación (IBMCC) for their technical assistance.

## CONFLICT OF INTEREST

The authors declare that they have no conflicts of interest.

## FUNDING

This work was mainly supported by a grant from the Fondo de Investigaciones Sanitarias (FIS) of the Spanish Ministry of Economy and Competitiveness and the European Regional Development Fund (ERDF) “Una manera de hacer Europa” [grant PI17/01895 to IGT and MSM.]; Junta de Castilla y León, Fondos FEDER [SA085U16 to JMHR]; Novartis grant; and by the Fundación “Jabones para Daniel”. JM Hernández-Sánchez was supported by a research grant from Fundación Española de Hematología y Hemoterapia (FEHH).

